# Diverse B-cell specific transcriptional contexts of the BCL2 oncogene in mouse models impacts pre-malignant development

**DOI:** 10.1101/2021.10.28.466164

**Authors:** Lina Zawil, Tiffany Marchiol, Baptiste Brauge, Alexis Saint-Amand, Claire Carrion, Elise Dessauge, Christelle Oblet, Sandrine Le Noir, Frédéric Mourcin, Florence Jouan, Mylène Brousse, Paco Derouault, Mehdi Alizadeh, Yolla El Makhour, Céline Monvoisin, Simon Léonard, Stéphanie Durand-Panteix, Karin Tarte, Michel Cogné

## Abstract

Follicular lymphoma (FL) is the most common indolent form of non-Hodgkin lymphoma arising from malignant germinal center (GC) B-cells. The genetic hallmark that leads to the development of FL is the t(14:18) which occurs early in the bone marrow during B cell development, thereby placing the anti-apoptotic *BCL2* gene under the direct control of the transcriptional enhancers in 3’ of immunoglobulin heavy chain locus (IgH 3’RR) and leading to the constitutive expression of the BCL2 protein. To assess the impact of the BCL2 deregulation on B-cell fate and try to reproduce FL development in mice, two models were designed: the Igκ-BCL2 (Knock in of the BCL2 in the light chain Ig kappa locus) and the 3’RR-BCL2 (Transgene containing BCL2 and a micro-3’RR), both containing the full BCL2 promoter region.

## Introduction

Follicular lymphoma (FL) is the most frequent indolent non-Hodgkin’s lymphoma (NHL), representing about 25% of B-cell malignancies (Milpied et al. 2021). This germinal center (GC) B-cell malignancy results from the malignant and clonal accumulation of centrocytes and follows a multistep lymphomagenesis process evolving over decades before clinical manifestations. After a long premalignant phase, the clinical course is slow and associates with multiple relapses associated with increasing resistant to therapy. Over time, approximately 30% of FL cases transform into aggressive diffuse large B-cell lymphoma (DLBCL).

The FL genetic hallmark is the translocation t(14; 18) (q32, q21) found in 90% of FL cases and deregulating the anti-apoptotic factor BCL2 under the control of immunoglobulin (Ig) heavy chain promoter. A translocation less frequent in FL, t(3;14), involves BCL6 such, hereby indirectly deregulating BCL2 (Ruminy et al. 2006). Contrasting to MYC in Burkitt lymphoma, for which variant translocations onto Ig light chain loci are common, *BCL2* t(2;18) and t(18;22) translocations are rare in FL but found in 9% of chronic lymphocytic leukemia cases (Hillion et al. 1991; Lin et al. 2008). Beside translocations, chromosomal amplification of *BCL2* is also observed in DLBCL and in mantle cell lymphoma (Monni et al. 1997). The precise pattern of BCL2 deregulated expression is thus likely impacting the phenotype of *BCL2*-driven lymphoproliferative disorders. The t(14;18) translocation stands as an aberrant V(D)J recombination product, joining double-strand breaks (DSBs) induced by RAG both in the IgH JH region (at position 14q32) and close to *BCL2* oncogene (at position 18q21) and hereby imposing an IgH-like pattern on *BCL2* gene accessibility and transcription (Raghavan et al. 2004). The *BCL2* gene encodes a transmembrane mitochondrial protein with dual roles, both inhibiting apoptosis and cell cycle entry. In mature B cells, expression of the IgH-translocated *BCL2* is mostly under the control of the IgH transcriptional enhancers located at the 3’ end of the IgH locus, which are the main IgH locus drivers at mature stages (Pinaud et al. 2011). These enhancers notably ensure accessibility of the locus to class switch recombination (CSR), somatic hypermutation (SHM) and hyper-transcription in activated B cells or plasma cells (Cogné et al. 1994; Rouaud et al. 2013; Saintamand et al. 2015; Pinaud et al. 2011). The functional result of this translocation is the imbalance between cell survival and death at a stage where mature B cells are intensely exposed to affinity-based selection within the GC. Normal GC B cells are indeed prone to undergo cell death unless they are positively selected and induced to enter the memory B cell or the plasma cell compartments. (Mayer et al. 2017; Péron et al. 2012). BCL2 is normally repressed in the GC while apoptosis is promoted by proteins such as Bak, Bax and Bad. BCL2 deregulation yielded by t(14; 18) thus jeopardizes the GC B cell survival checkpoint and *BCL2* translocation stands as a driver genetic event initiating lymphomagenesis in FL.

Among naive B cells having left the BM and circulating in the periphery, those harboring the t(14;18) translocation display a selective advantage during the GC reaction allowing them to persist as atypical memory B cells already carrying some features of FL cells and considered as FL precursor cells. These premalignant cells are prone to re-enter the GC upon future Ag encounter, undergoing additional SHM and eventual additional oncogenic hits under the iterative exposure to AID activity (Milpied, Nadel, and Roulland 2015; Tellier et al. 2014; Huet, Sujobert, and Salles 2018). Although an early and crucial anomaly, the t(14;18) by itself is not sufficient for FL development since the prevalence of FL does not exceed 0.03%. Among the most frequent additional hits, alterations of histone/chromatin modifying enzymes, including KMT2D, and CREBBP, are collectively found in almost 100% of FL cases whereas mutations of HVEM/TNFRSF14 or introduction of N-glycosylation sites within Ig variable regions have been shown to affect the crosstalk between tumor B cells and their surrounding microenvironment (Boice et al. 2016; Amin et al. 2015; Huet, Sujobert, and Salles 2018).

Exploring the mechanisms of such a complex disease requires to study whole living organisms and has prompted the generation of several models in mice. Initial transgenic mouse models expressing BCL2 driven by the Eμ enhancer have mostly resulted in polyclonal expansion of all B-cell compartments overexpressing BCL2, from progenitors to plasma cells (McDonnell et al. 1989; Strasser et al. 1991). Lymphoma development was reported mostly when the BCL2 deregulation was associated with other spontaneously selected or experimentally enforced genetic anomalies, notably involving Myc (Strasser et al. 1990; McDonnell and Korsmeyer 1991). However, lymphoproliferation observed in such conditions rather involved immature B cells and did not represent models for post-GC low-grade human lymphoma. Strikingly, the most widely used transgenic mice considered as a pertinent model of human FL and based on the tumorigenic potential of BCL2 have been the vavP-*BCL2* transgenics, although they broadly overexpress BCL2 in all hematopoietic lineages, including T-cells and thus cannot recapitulate the natural story of human FL where BCL2 overexpression is strictly restricted to mature B cells (Ogilvy et al. 1999; Egle et al. 2004).

Therefore, in an aim to better understand the complex development of FL and the impact of *BCL2* deregulation, our current study explores two new mouse models, designed either for pan-B cell expression through a knock-in of *BCL2* in the Igκ locus, or for a specific targeting of activated B-cells with a *BCL2* transgene driven by the IgH 3’RR enhancers. Since these elements are major drivers of Ig gene remodeling in the GC, the latter model is expected to more specifically mimic the BCL2 deregulation associated with FL. The *BCL2* promoter region has a characteristic structure comprising the P1 and P2 promoters, with P1 dominating in healthy lymphocytes, while a shift from P1 to P2 is observed in FL. Contrary to previous *BCL2* transgenes in the literature, the full human *BCL2* promoter region was thus included in our BCL2 gene cassettes. Indeed, this region includes transcription factor-binding sites, notably for the repressor BCL6, that limits *BCL2* expression in normal GC cells (Saito et al. 2009). The design of our constructs should thus allow to reproduce in mice the deregulation or the mutations of the BCL2 promoter region documented in patients. On their own, both models develop B-cell expansion which differed in terms of stage-specificity according to the context of *BCL2* deregulation, 3’RR-*BCL2* mice showing the most “GC-restricted” phenotype. These models are pertinent for studying a pre-FL stage in young mice, notably showing the impact of GC B-cell expansion on the TFH counterparts. In the absence of other genetic abnormality, neither of these models however develop FL-like lymphoma at late ages, but rather plasma cell tumors with mutated Ig genes corresponding to post-GC cells.

## Materials and methods

### 1 Cell lines and mouse models

DoHH2, SU-DHL-4, SU-DHL-6 and OCI-Ly3 cell lines (Germinal center or activated B cell type Diffuse large B cell lymphoma (DLBCL)) were grown in RPMI 1640 medium supplemented with 10% Fetal calf serum (Dominique Dutscher, Catalog number: S1810-500) at 37°C with 5% CO2 atmosphere).

All in vivo experiments were performed in accordance with animal ethical rules, and all protocols were authorized by the French Ministry of Research according to European Union regulations (APAFiS 13900). All mice were bred in a specific and opportunistic free (SOPF) animal facility.

Two new mouse models were designed for this study (Fig 1A):

**Figure 1:**
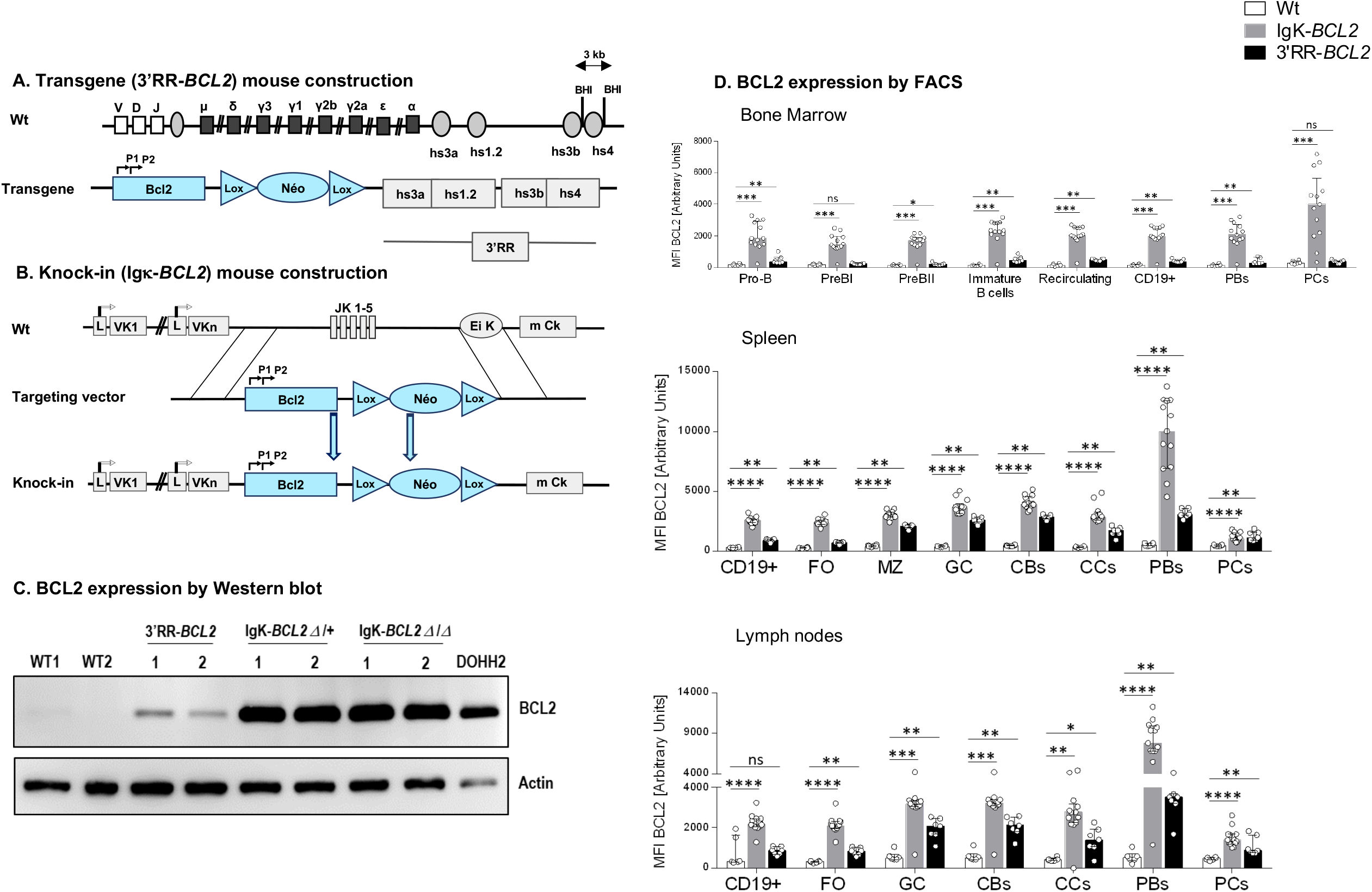
Mouse model construction schematics for (A) 3’RR-BCL2 transgenics and (B) Igκ-BCL2 knock-in mice. (C) Western-blot evaluation of human BCL2 protein expression in the spleen of WT (negative control), 3’RR-BCL2, Igκ-BCL2 Δ/+, Igκ-BCL2 Δ/Δ mice, and in the DoHH2 human cell line (positive control) (D) Human BCL2 expression in both mouse models, using the mean fluorescent intensity (MFI) obtained by flow cytometry from bone marrow, spleen and mesenteric lymph nodes B-cells from unimmunized 3’RR-*BCL2* and Igκ-*BCL2* Δ/+ only.

1. The “3’RR-*BCL2”* model, with random integration of a transgene which contains a human *BCL2* gene cassette driven by the *BCL2* P1/P2 promoter region, under the control of 3’ IgH superenhancer (a “micro IgH 3’RR” combining the various enhancer elements from the Ig heavy chain locus 3’ regulatory region)
2. The “Igκ-*BCL2”* model which is a deletion (Δ) of the Jκ region, replaced with the same BCL2 cassette [P1/P2 promoter *BCL2* gene] described above, thus as a knock-in in the mouse Igκ light chain locus, immediately upstream of the Eκ enhancer.

Both of our models thus contain the full promoter region of *BCL2*, including its two alternative promoters P1 and P2, all in a mixed B6;129S genetic background.

### 2 Flow cytometry analysis of lymphoid compartment

Single cell suspensions from the Bone marrow (BM), and secondary lymphoid organs (spleen and mesenteric lymph nodes) were taken from mice that were either resting (unimmunized) or immunized with 3 iterative monthly SRBC injections. These cell suspensions were then labelled using various extracellular antibodies designed to see the different early and late B cell populations as well as the different T cell populations in each of the organs mentioned above. Following the extracellular staining step, cells were fixed and permeabilized using the eBioscience FOXP3/transcription staining buffer set (Invitrogen, Reference: 00-5523-00) and then stained for the intracellular proteins. The detailed list of antibodies used in each of these panels is summarized in supplementary Table 1. Cells were then analyzed by flow cytometry using an LSR FORTESSA cytometer (Beckton Dickinson) and data analysis was done using the <Flow Logic> system (the precise gating strategies for the different panels are listed in supplementary Figure 2A-C). Percentages were determined (for the different B cell sub-populations among CD19+ cells and for the different T cell sub-populations among CD4+ cells) as well as absolute cell numbers only in immunized mice (calculated based on the measured cell counts and lymphocyte percentages in the organ analyzed).

The mice used for evaluating lymphoid compartments represented 2 cohorts:

1. Resting cohort: this cohort comprised of 6 wildtype, 13 hemizygous Igκ-*BCL2* Δ/+ (Igκ-*BCL2* Δ/+) and 7 3’RR-*BCL2* mice were analyzed to carry out full characterizations of our mice at the resting state.
2. Iteratively immunized cohort: 3 groups of 13 wildtype, 7 Igκ-*BCL2* Δ/+ and 10 3’RR-*BCL2* mice were immunized with the following protocol: at 3 months of age, mice were immunized intra-peritoneally with 200 μl of Sheep Red Blood Cells (SRBC) iteratively over 3 consecutive months and were sacrificed one month after the third immunization.

### 3 Western Blots

For detection of human BCL2, we used a cohort of 2 wildtype mice (as a negative control), 2 3’RR-*BCL2*, 2 homozygous Igκ-*BCL2* (Igκ-BCL2 Δ/Δ), 2 hemizygous Igκ-*BCL2* (Igκ-BCL2 Δ/+) and 1 DoHH2 cell line (as a positive control).

10 μg of total proteins were extracted (using 2X Laemmli buffer, Biorad, Catalog number # 161-0737, Composition: 65.8 mM Tris-HCl PH 6.8, 26.3% (w/v) glycerol, 2.1% SDS, 0.01% bromophenol blue) and denatured / reduced using Δ-mercaptoethanol (2.5% final) at 95 °c for 5 mins.

After electrophoresis on a 12% polyacrylamide gel (Biorad), proteins were transferred onto PVDF membranes (GE Healthcare). The human BCL2 protein was detected using the same mouse anti-human BCL2 antibody we use for flow cytometry followed by an HRP-linked anti-mouse Ig antibody (eBioscience, #18-8817-30).

Actin was used as a housekeeping protein (Rabbit anti-Actin antibody (Sigma-Aldrich A2066) followed by Goat anti-Rabbit-HRP (Southern Biotech 4050-05)).

Membranes were developed by enhanced chemiluminescence for high sensitivity detection system according to the manufacturer’s instructions (Biorad).

### 4 Proliferation Test

A separate cohort of 6 wild type (WT), 4 Igκ-*BCL2* Δ/+ hemizygous mice and 5 3’RR-*BCL2* mice were injected intraperitoneally with 200 μl of SRBC at day 0, followed by another intraperitoneal injection with 200 μl of 5-ethynyl-2′-deoxyuridine (EdU) at Day 6 and they were sacrificed at day 7.

Single cell suspensions were taken from the bone marrow, spleen and mesenteric lymph nodes and stained using the same panel described above but modified to be able to see the EDU+ cells. This was done following the “Click-iT EdU Flow Cytometry Assay Kit protocol” (Molecular probes by Life technologies, catalog numbers C10419, C10420). Cells were then analyzed by flow cytometry using the BD LSR FORTESSA and data analysis was done using the <Flow Logic> system.

### 5 Quantification of Antibody affinity

We followed the indirect enzyme-linked immunosorbent assay (ELISA) described by Zhang et al. to quantify the affinity of the produced antibodies (Notably IgG) in both of our mouse models compared to wild type.

Mice were injected with Ovalbumin at 1ng/ml and Addavax (equal volume for both) at Day 0 and Day 14. Sera was taken at different time points to follow up the secretion of IgM and IgG (Day 0, 7, 14, 21 and 28). Only sera of day 28 were eventually studied. Quantification of total IgG in the serum was first done followed by the ELISA based affinity study.

96-well plates were coated with Ovalbumin at 2 μg/ml. In parallel, sera of Day 28 for each mouse model (at a fixed concentration of 5×10^−10^ M for IgG) was incubated overnight with Ovalbumin at different concentrations but always in large excess compared to IgG (4×10^−7^ to 6.25×10^−9^ M). The next day, the mix of serum IgG / Ovalbumin was incubated with the plate previously coated with Ovalbumin. Then, the alkaline phosphatase conjugated secondary antibody (anti-Mouse IgG) was added. After 45 minutes of incubation, P-Nitrophenyl phosphate (1 mg/ml) was added and alkaline-phosphatase activity was blocked with 3M Sodium hydroxide (NaOH) and the optical density was measured at 405 nm using a Multiskan FC photometer.

Finally, the dissociation constant (K_D_) was evaluated by measuring the slope of the linear dependence A_0_ / (A_0_ – A) upon 1/a_0_ (Zhang et al. 2012).

### 6 Next-generation sequencing for repertoire analysis

We performed repertoire sequencing analysis on RNA (500ng - 1.5mg) extracted from:

1. Primary cell suspensions of tumoral tissue, spleen and mesenteric LNs of 5 Igκ-*BCL2* Δ/Δ tumoral mice, 3 Igκ-*BCL2* Δ/+ tumoral mice and only one 3’RR-*BCL2* tumoral mouse
2. Primary cell suspensions of bone marrow, spleen and mesenteric LNs of 5 Igκ-*BCL2* Δ/+ mice, 5 3’RR-*BCL2* mice and 9 wildtype mice from the iteratively immunized cohort.

Sequencing was done using the strategy described by Li et al. developed for T-cell repertoire diversity and clonotype. These experiments used a new generation methodology, which combines 5’ RACE PCR; sequencing; and, for analysis, the international ImMunoGeneTics information system (IMGT), IMGT/HighV-QUEST Web portal, and IMGT-ONTOLOGY concepts. Briefly, we amplified transcripts with 5’ RACE PCR using a reverse primer hybridizing within the Mu (μ) and gamma (γ). Sequencing adapter sequences were thus added by primer extension, and resulting amplicons were sequenced on a GS FLX 1 Sequencing system (Roche, Pleasanton, CA). Repertoire was done using IMGT/High-V-Quest and associated RStudio package scripts; associated tools are available on the IMGT Web site.

### 7 Single cell sequencing and analysis

Three sets of 10-day splenic B cells were magnetically sorted (StemCell B cell negative selection kit) from mice immunized once with SRBC (EasySep™ Mouse B Cell Isolation Kit (Catalog # 19854)). Single cells were captured and barcoded using the 10X 3’ sequencing kit (Chromium Next Gem Single cell 3’ reagent kit v3.1, 10X Genomics), and libraries were prepared following the manufacturer’s instruction. Libraries were run using 2×75 paired end reads on the HiSeq4000 Illumina sequencer. Raw data were successively processed and analyzed with the 10X Cell Ranger (10XGenomics) and Seurat (v3.2.3) package (Stuart et al. 2019). Mean reads per cell was 40,766 for the WT sample, 66,971 for the 3’RR-BCL2 sample and 102,596 for the Igκ-BCL2 sample. Median genes per cell over 3 samples varied from 1767 to 1869. Cells expressing less than 800 or more than 4500 genes or with more than 20000 umi counts were filtered out. Cells with the frequency of mitochondrial genes more than 8% or with the frequency of ribosome genes less than 10% were also removed from the analysis. Contaminating T-cells and myeloid cells were detected based on canonical markers (Cd3 genes and C1q genes) and filtered out. Gene counts were normalized using the SCTransform package (v0.3.1) with a second non-regularized linear regression applied to percentage of mitochondrial and ribosomal genes (Hafemeister and Satija 2019). Canonical correlation analysis was used to integrate data from different batches by running the following steps as implemented in Seurat package (Butler et al. 2018): Selection of integration features; Removal of immunoglobulin genes from the features used for integration; Preparation for integration with the PrepSCTIntegration function. Data integration was then performed using the FindIntegrationAnchors and IntegrateData functions based on the first 30 correlation components. PCA analysis was performed on the integrated dataset and the first 12 principal components were used for UMAP computation and clustering (using 30 nearest neighbors and a resolution of 0.2). Cluster 1 was subclustered by executing the same steps previously described (PCA analysis on the subset and clustering computation with the first 10 principal components, 15 nearest neighbors and a resolution of 0.2). Using lognormalized expression values, marker genes for each cluster as well as differentially expressed genes between conditions were inferred by the Wilcoxon test as implemented in the FindAllMarkers function. Cell cycle score and classification were calculated by CellCycleScoring function based on the expression of G2/M and S phase markers (Kowalczyk et al. 2015).

### 8 Tumor Follow up

A cohort of 12 Igκ-*BCL2* Δ/Δ, 7 Igκ-*BCL2* Δ/+ and 8 3’RR-*BCL2* mice were monitored for the spontaneous development of tumors with time.

### 9 Study of BCL2 promoter mutations

For this study, DNA was phenol/chloroform extracted from:

1. The various cell lines listed in section 1 (DoHH2, SU-DHL-4, SU-DHL-6 and OCI-Ly3) used as positive controls for BCL2 promoter mutations.
2. Total spleen from iteratively immunized mice (4 Igκ-*BCL2* Δ/+ and 4 3’RR-*BCL2*).
3. Class switched versus non- class switched B cells from additional groups of 3 months old immunized Igκ-*BCL2* Δ/+ (3 mice) and 3 3’RR-*BCL2* mice (3 mice). These mice were induced only once with SRBC and sacrificed at day7. Single cell suspensions from the spleen were stained with CD19 coupled to BV510 (BD Biosciences, Clone: 1D3) and IgM coupled to FITC (Southern Biotech, Cat number: 1020-02) antibodies. Cells were then sorted using the BD Aria III cell sorter to obtain class switched B cells (CD19^+^/IgM^-^) and non-class switched B cells (CD19^+^/IgM^+^).
4. From mice tumors:
  - Tumoral tissues of 3 Igκ-*BCL2* Δ/+ Mice
  - Mesenteric LNs of one Igκ-*BCL2*, used as positive controls Δ/+ and one 3’RR-*BCL2* tumoral mice
  - Total spleen of two 3’RR-*BCL2* tumoral mice

The full promoter region of *BCL2* was then amplified by polymerase chain reaction (PCR) using specific forward (5’ TGAATGAACCGTGTGACGTTACGC 3’) and reverse (5’ CTCAGCCCAGACTCACATCA 3’) primers. The amplification of the PCR product (2,184 bp long) is verified by gel electrophoresis using agarose 2% gel in TBE. The amplified PCR product is then purified from gel using the “Nucleospin Gel and PCR Clean-up” kit (Macherey-Nagel, Reference: 740609.250).

Next-generation sequencing was performed according to the user guide of the Ion Xpress Plus gDNA Fragment Library Preparation (Life Technologies catalog no. 4471269;) from amplified products (1 μg), and libraries were sequenced on an Ion Proton System. Two negative controls are mandatory for each library run. In this study, we our negative controls comprised of genomic DNA samples extracted from the tails of an Igκ-*BCL2* Δ/+ and a 3’RR-*BCL2* mouse respectively. Analysis was done using Deminer software developed by our laboratory in order to subtract the background level of mutations observed on the same sequence in a control experiment (Martin et al. 2018).

## Results

### Generation of mice carrying B-cell specific *BCL2* deregulation

Mice hemizygous for any of our 2 *BCL2* transgenic models were studied and produced high amounts of human BCL2 of normal size in B-lineage specific manner (Fig 1A-D). Both revealed overexpression of the human BCL2 protein in spleen follicular and GC B cells with a stronger BCL2 expression in the Igκ-*BCL2* than in the 3’RR-*BCL2* mice (Fig 1D. In addition, BCL2 expression patterns along B cell differentiation stages strongly differed (Fig 1D). The Igκ-locus driven deregulation consisted in very high and stable BCL2 overexpression throughout B-cell differentiation in the bone marrow from B cell progenitors to recirculating mature B cells, plasmablasts having migrated to the bone marrow and plasma cells. This architecture of the transgene also yielded a constitutively high expression in peripheral spleen and lymph node lymphoid tissues, homogeneously affecting all B-cell compartments, but strongly culminating in plasmablasts.

In striking contrast, the IgH 3’RR-BCL2 drove low BCL2 expression in bone marrow B cell progenitors and plasma cells. In spleen and lymph nodes, it provided the highest expression in GC B cells, either centrocytes or centroblasts, with expression remaining high in plasmablasts but rapidly falling down in plasma cells.

### Impact of *BCL2* deregulation on B and T cell differentiation

We analyzed early B cell compartments by flow cytometry in non-immunized mice bred in an SOPF facility, comparing mutant mice to WT controls. Prior to any immunization, the percentage of bone marrow CD19+ cells was similar to that in WT controls and showed no amplification of any progenitor compartment, even with rather decreased percentages of pro-B and pre-B cells in Igκ-BCL2 mice (**Fig 2A**). By contrast, bone marrow plasmablasts and PCs were constitutively increased in the Igκ-BCL2 mice.

**Figure 2:**
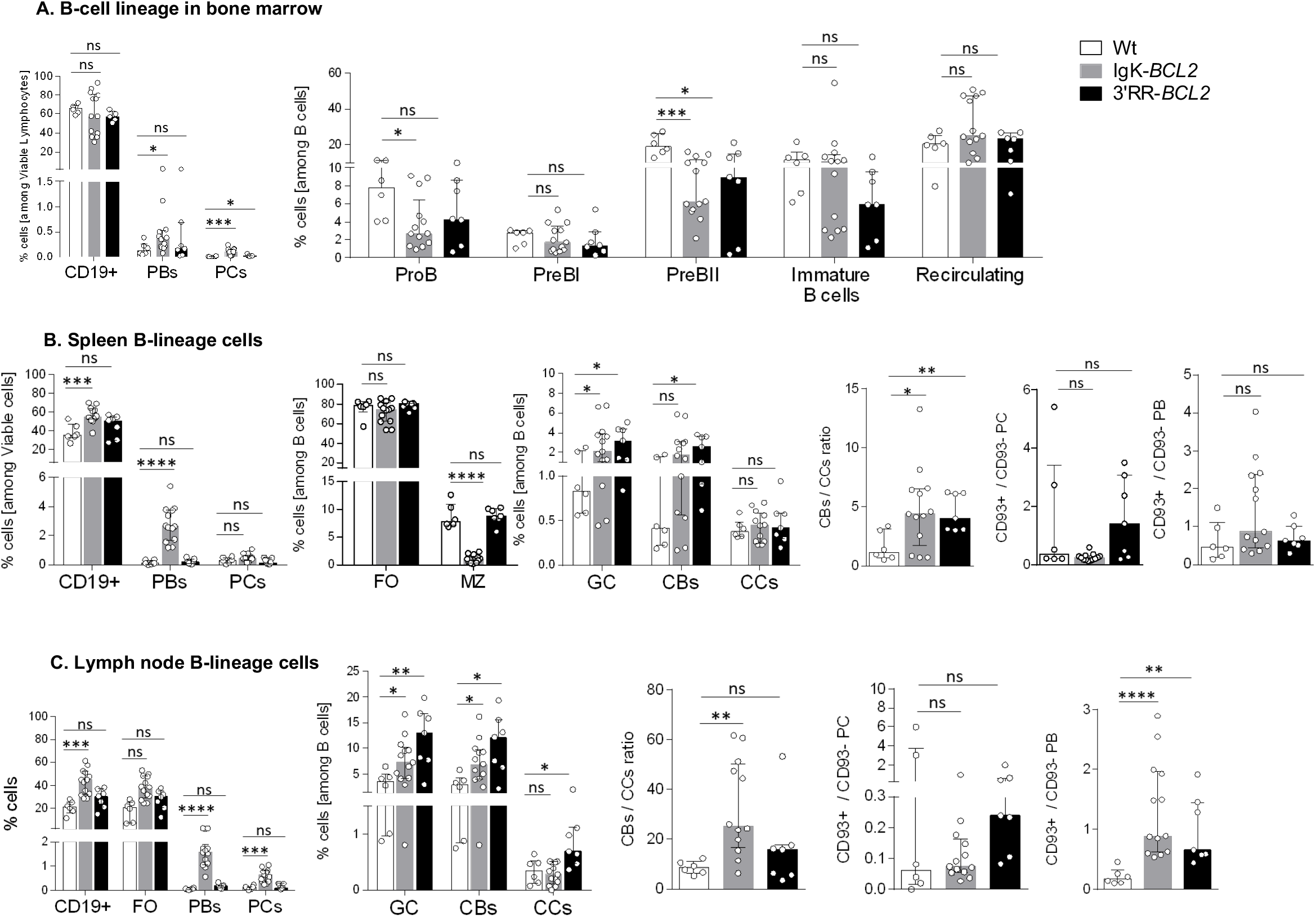
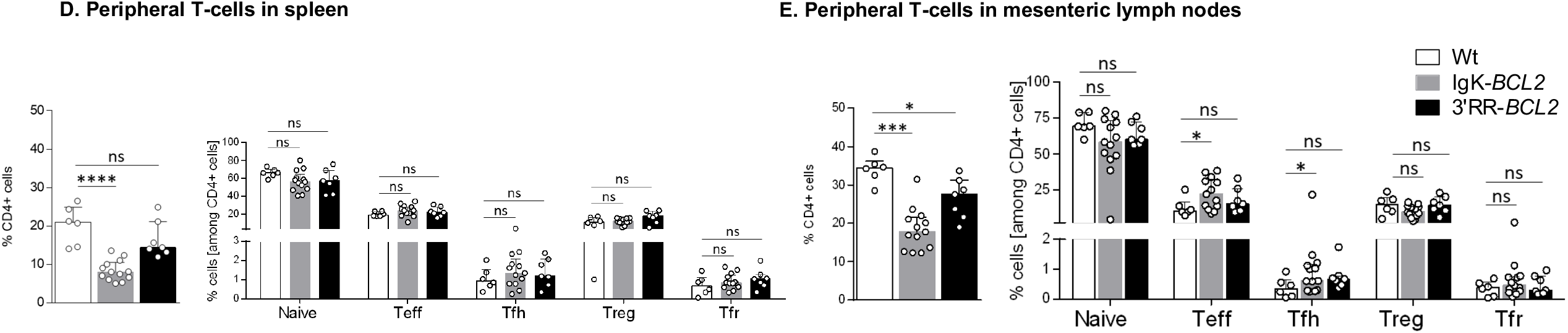
Flow cytometry data showing the impact of BCL2 deregulation alone, in both mouse models compared to WT mice at the resting non-immunized stage, on the different B cell compartments, in bone marrow (A), spleen (B) and mesenteric lymph nodes (C), as well as on the T cell compartment (D). Percentages of the different cell compartments are presented as medians with interquartile range and the statistical significance was determined by the Mann-Whitney *U* test. Ratios for the centroblasts to centrocytes, as well as CD93+ to CD93-plasma cell and plasmablastic cell populations are also presented. The full staining panel of both B and T cells in each of the organs is summarized as supplementary Table 1.

In the periphery of resting unimmunized Igκ-*BCL2* mice, CD19+ cells were globally more abundant than in WT controls in both lymph nodes and spleen, while 3’RR-*BCL2* mice inconstantly showed such an increase. The difference between both strains was still more striking for plasmablasts and plasma cells, which were in normal amount in 3’RR-*BCL2* mice while they increased in number by up to ten-fold in Igκ-*BCL2* (**Fig 2B and 2C**). By contrast, marginal zone (MZ) cells were decreased in Igκ-*BCL2* mice. The ratio of CD93+CD138+ *vs* CD93-CD138+ plasmablasts was rather increased in both strains, significantly in lymph nodes, suggesting increased amount of recently differentiating plasmablasts (Chevrier et al. 2009). In PCs, for which CD93 expression marks long-lived PCs having arisen from T-dependent responses, there was a tendency to an increased ratio of such cells in the 3’RR-BCL2 mice, but not reaching significance (Chevrier et al. 2009).

Regarding the GC B-cell compartments in spleen and lymph nodes (corresponding to background GCs in non-immunized mice), both strains showed a constitutive increase of both centroblasts and centrocytes and an increased ratio of centroblasts *vs* centrocytes. Relatively to expanded B cell populations, the percentage of helper T cells appeared significant decreased in the periphery of Igκ-*BCL2* mice only (**Fig 2D**), with a tendency for both BCL2 strains to have less naïve T cells but slightly increased effector (Teff) and follicular helper T (Tfh) cell populations, only showing statistical significance in the mesenteric LNs of the Igκ-*BCL2* mice (**Fig 2E**).

### Lymphoid compartments in immunized mice

Human lymphoid malignancies are often correlated with past chronic activation by viral Ag or auto-Ag, and we thus explored the behavior of BCL2 transgenic mice in conditions of 3 consecutive B-cell stimulations with the particulate Ag sheep red blood cells

(SRBCs). Such a repeated stimulation resulted in a strong global increase of the B-cell compartment in all lymphoid tissues from Igκ-*BCL2* mice, but not 3’RR-*BCL2* mice (**Fig 3**). The Igκ-BCL2 also confirmed an MZ B-cell decrease.

**Figure 3:**
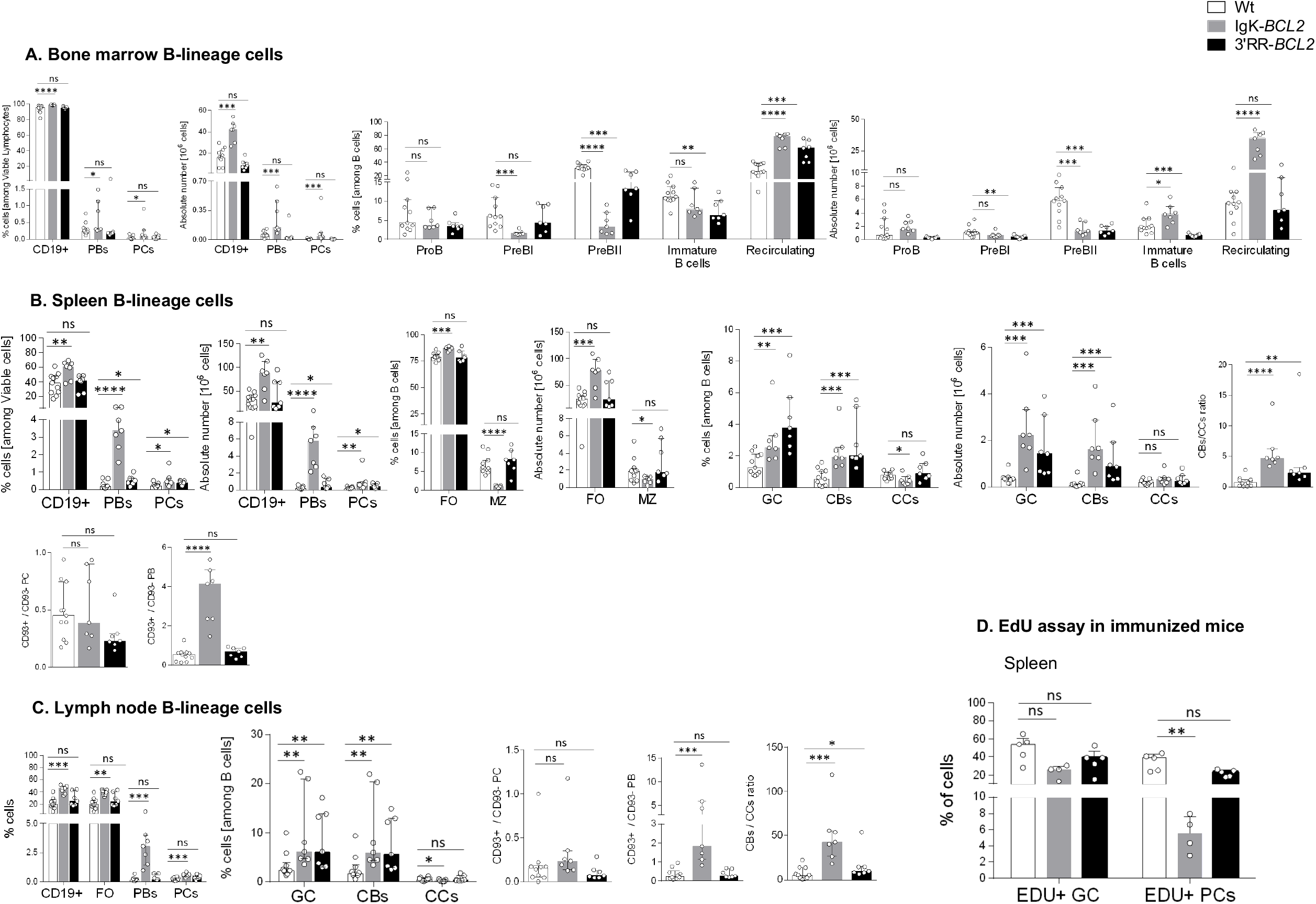
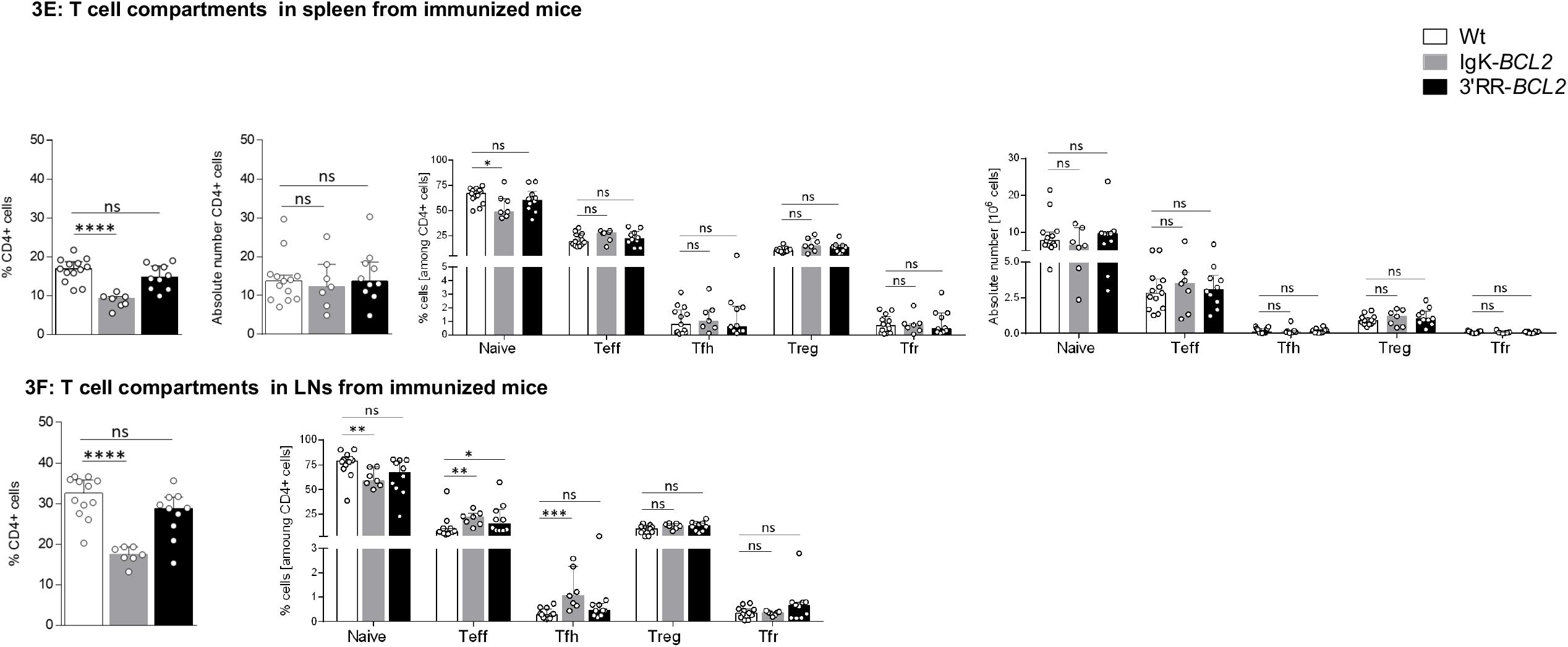
Flow cytometry data showing the impact of the BCL2 deregulation accompanied with 3 iterative intra-peritoneal SRBC injections (on a monthly basis) on the different B cell populations, in both BCL2 mouse models compared to WT, in the early stages of development in the bone marrow (A) and late ones in the secondary lymphoid organs, the spleen and mesenteric lymph nodes (B and C respectively). The ratios of centroblasts to centrocytes as well as CD93+ to CD93-plasma cells and plasmablasts are also indicated. Percentages of the different cell populations as well as absolute cell numbers (calculated based on the percentages obtained by flow cytometry and the total percentage of lymphocytes in each corresponding organ) are presented as medians with interquartile range and the statistical significance was determined using the Mann-Whitney *U* test. The full staining panel for each organ as well as the gating strategy used are summarized in supplementary Table 1 and supplementary Figure 1A-C. (D) Flow cytometry data showing the expression (or not) of EDU in splenocytes, 24 hours post its intraperitoneal injection into 5 WT, 4 Igκ-*BCL2* Δ/+ and 5 3’RR-*BCL2* mice (and 7 days after SRBC immunization). The percentages of EDU expressing plasma cells (EDU+ PC) and GC B cells (EDU+ GC) are presented as medians with interquartile range. Statistical significances were realized using Mann-Whitney *U* test. The EDU used for the staining panel was coupled to FITC (A488) along with markers specific for the plasma cell (CD138 BV786), GC B cells (GL7 coupled to APC and CD38 coupled to APC-R700) and cell viability (FVS 780). (E) Flow cytometry data showing the impact of the BCL2 deregulation accompanied with 3 iterative intra-peritoneal SRBC injections (on a monthly basis) on the different T cell populations, in both BCL2 mouse models compared to WT, in the periphery (spleen and LNs). Both percentages and absolute cell numbers each T cell population are presented as medians with interquartile range and the statistical significance is realized via the Mann-Whitney *U* test.

In bone marrow (**Fig 3A**), the B-cell increase corresponded to recirculating B-cells, whereas all B-cell progenitor compartments appeared to be decreased. Percentages and absolute number of bone marrow plasmablasts and plasma cells were also significantly increased in Igκ-BCL2 mice.

In spleen and lymph nodes from Igκ-*BCL2* mice (Fig 3B-C), the global increase of CD19+ cells also associated with a strong persisting increase of plasmablasts and to a lower extent of plasma cells. Plasma cells quantified after immunization did not show the abovementioned increased ratio of CD93+ cells and not yet express the markers of LLPCs. By contrast for the burst of plasmablasts seen in Igκ-BCL2 mice, a strongly increased ratio of CD93+CD138+ plasmabasts was seen in spleen, corresponding to a high amount a recently differentiated cells (Chevrier et al. 2009).

In order to evaluate whether the plasmacytosis in Igκ-BCL2 mice rather involves the ongoing entry of B-cells into the PC stage or the accumulation of long-lived PCs, we measured BrDU incorporation into PCs and GC B-cells one week after immunization (**Fig 3D**). This experiment revealed a decreased ratio of BrDU+/ BrDU PCs, indicating that accumulation of long-lived PCs predominantly contributed to the peripheral plasmacytosis. Parallel evaluation of BrdU incorporation in GC B cells showed a preserved ratio of recently divided *vs* lately divided cells.

By contrast to Igκ-*BCL2* mice, 3’RR-*BCL2* mice showed a more specific relative increase of GC B-cells, involving both centrocytes and centroblasts, together with a higher ratio of centroblasts *vs* centrocytes, suggesting that BCL2 expression in these mice mostly impacted B-cell survival at the centroblast stage.

Anomalies of T-cell compartments after repetitive immunization remained similar to those in resting mice, with a significant decrease of the peripheral helper T cell population in the Igκ-*BCL2* mice, together with increased Teff and TFH cell populations in the LNs of both models (**Fig3E-F**).

### Antibody affinity

In parallel, we wanted to assess the ability of these mice to mount an efficient immune response against a T-dependent Ag despite their altered GC regulation. We quantified the affinity of anti-Ovalbumin IgG antibodies using the ELISA-based protocol. Analysis of the obtained dissociation constant (K_D_) showed that both models are able to produce large amount of high affinity antibodies (the higher the affinity of the produced antibody the lower the K_D_). With an ELISA-based evaluation of antibody affinity, both of our models produced high-affinity circulating IgG with dissociation constant similar to WT (or even with a tendency towards a lower value) (Fig 4). This finding is reminiscent from observations previously done for Eμ-BCL2 mice, in which the selection of Ag-specific B-cells appeared to maintained (Smith et al. 2000).

**Figure 4:**
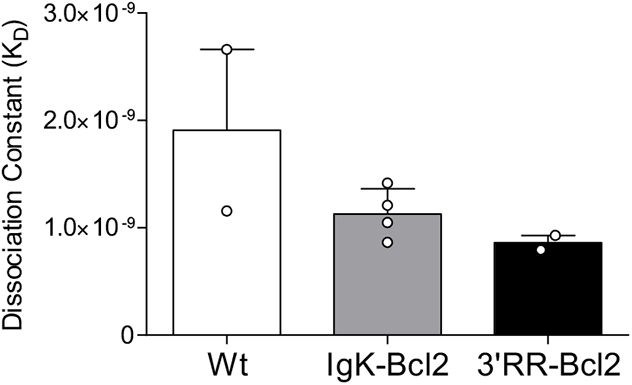
(A) Histogram showing the dissociation constant (K_D_) of IgG to Ovalbumin, obtained by an ELISA-based assay (described in materials and methods), in the sera of both BCL2 mouse models compared to WT after 45 minutes of incubation with the secondary antibody. A lower KD value indicated higher IgG affinity against Ovalbumin. K_D_ is reported here as medians with interquartile range and no statistical significance was determined due to the limited number of mice.

### Single cell transcriptome analysis

In order to appreciate the impact of BCL2 deregulation on the B-cell transcriptome more precisely than by following surface expression of a limited set of membrane markers, we carried a single cell analysis of B-lineage splenocytes 10 days after immunization. This analysis allowed to identify 12 clusters and sub-clusters. Top differentially expressed genes are indicated in Supplementary Figure 2. From their transcriptional profile, clusters were identified as transitional cells, recirculating resting B-cells, extrafollicular and interfollicular cells, IFN-activated cells, marginal zone, pre-centroblastic, centroblasts, early centrocytes (AID+ BCL6+), late centrocytes (AID-, BCL6+), preplasmablasts, plasmablasts and plasma cells (**Fig 5A**). This analysis confirmed and extended the observations made by flow cytometry, but with much better precision. All compartments upstream of GC formation were rather decreased in BCL2 transgenics (from transitional cells to resting, extrafollicular, IFN-activated, marginal zone and pre-centroblastic cells⃛ In the Igκ-BCL2 model, the compartments amplified began with centrocytes but culminated with plasmablasts and plasma cells. By contrast in the 3’RR-BCL2 model, the amplified subclusters were more “GC-focused”, including centroblasts, centrocytes, late centrocytes and plasmablasts (**Fig 5B**). This analysis also allowed to monitor the mean expression level of a number of highly expressed genes comparatively between WT, Igκ-BCL2 and 3’RR-BCL2 mice. With more precision than in flow cytometric measurements, this analysis showed a broad overexpression of BCL2 in all subclusters in the Igκ-BCL2 model, compared with a lower and much more specific expression in 3’RR-BCL2 B cells, focused on centroblasts, centrocytes and plasma cells (**Fig 5C**).

**Figure 5:**
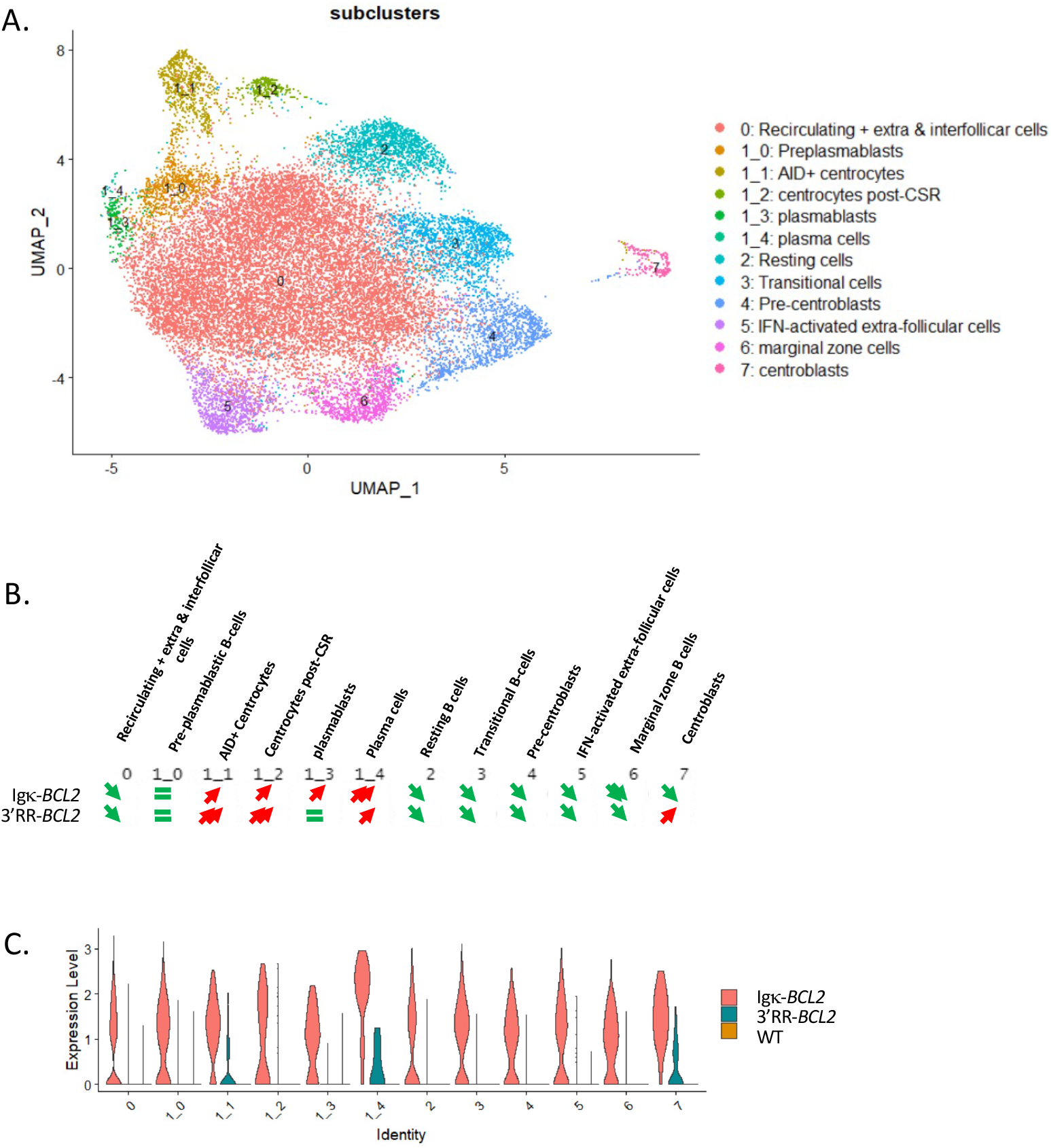
(A) UMAP of the 10X single cell analysis of spleen B-cells 10 days after SRBC immunization, split into 12 clusters, (as defined in Supplementary Figure 2). (B) Quantitative variations of cell numbers in the various clusters in Igκ-*BCL2* and 3’RR-*BCL2* mice compared to WT. (C) Human BCL2 expression level in the various cell clusters in Igκ-BCL2 and 3’RR-*BCL2*.

### B-cell diversity in immunized mice

Repertoire of immunized mice only showed minor changes by comparison to normal mice (**Fig 6**), without significant increase of the Gini index in any of the Ig transcript category analyzed (μ or γ IgH transcripts from spleen or lymph nodes), thus indicating at this stage that no clonal amplification was detectable but that normally diversified repertoires were expressed after immunization. Usage of VH subgroups associated with IgH μ or IgH γ transcripts showed no significant variation compared to WT.

**Figure 6:**
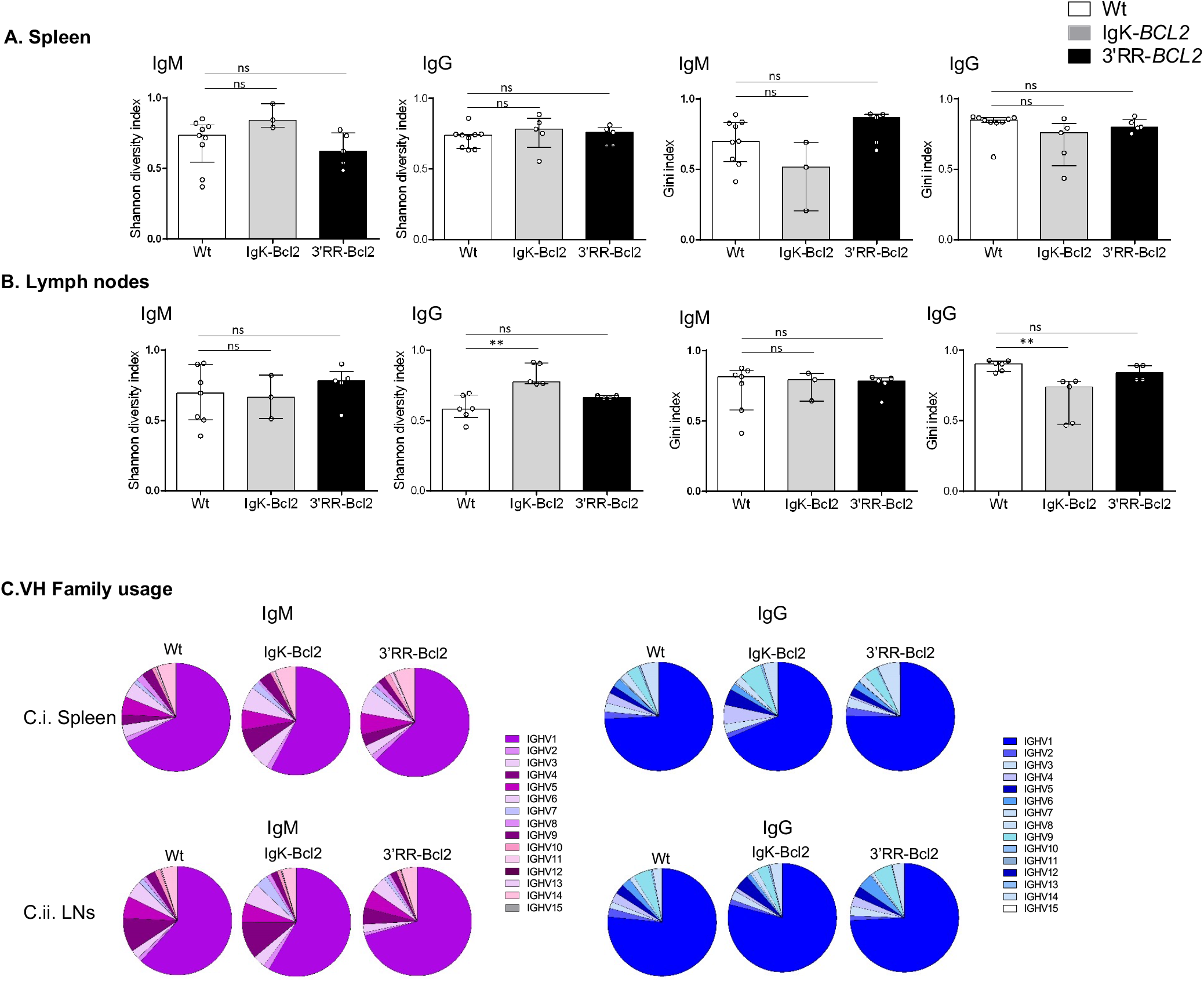
Histogram showing the Shannon diversity index and the Gini index of both IgM and IgG repertoires in the spleen (A) and mesenteric LNs (B) of both BCL2 mouse models compared to WT. Both indices are presented as medians with interquartile range and statistical significances are determined by Mann-Whitney *U* test. (C) Pie charts showing the mean percentage of each VH gene family usage of both IgM (in purple) and IgG (in blue) in the spleen and mesenteric LNs of both BCL2 mouse models compared to WT.

### Immunohistochemical analysis and tumor development in *BCL2* transgenic mice

We allowed cohorts of 40 mice for each genotype to grow older, in order to monitor the potential onset of spontaneous tumors. Aggressive tumors indeed started to appear after around 25 weeks of age in about 15% of mice with either configuration of the BCL2 (**Table 1** and **Fig 7A)**.

**Table 1:** Table describing in the mice that developed tumors in both BCL2 models. This cohort comprised of 19 Igκ-BCL2 (total) and 8 3’RR-BCL2 mice was allowed to grow older and these tumors were monitored for tumor development over time. For each mouse, the age at which they developed the tumor (or pre-tumoral symptoms) along with detailed description of the location and the phenotype of each tumor is listed.

| Genotype | Number | Age (weeks) | Ascitis | Tumor | Organs affected / Tumor description | Other observations |
| --- | --- | --- | --- | --- | --- | --- |
| IgK-Bcl2 Δ/Δ | KI- 1 | 85 | ✓ | ✓ | Intestine, Mesenteric LNs, Cervical LNs, Splenomegaly (165 mg) | / |
|  | KI- 2 | 50 | X | ✓ | Intestine, Mesenteric LNs, Cervical LNs, Splenomegaly (650 mg) | / |
|  | KI- 3 | 71 | ✓ | ✓ | Intestine, Stomach, Splenomegaly (395 mg) | / |
|  | KI- 4 | 90 | X | ✓ | Intestine, LNs | occlusion |
|  | KI- 5 | 71 | X | ✓ | Mesenteric LNs, Liver, Splenomegaly (1580 mg) | / |
|  | KI- 6 | 54 | ✓ | ✓ | Stomach, Intestine , Mesenteric LNs, Liver, Pancreas, Splenomegaly (305 mg) | / |
|  | KI- 7 | 54 | ✓ | ✓ | Kidneys, LN atrophy, Splenomegaly (950 mg) | / |
|  | KI- 8 | 105 | X | ✓ | Mesentery, Liver, Kidneys, Splenomegaly | / |
|  | KI- 9 | 109 | ✓ | ✓ | Liver, Splenomegaly (397 mg) | / |
|  | KI- 10 | 70 | X | ✓ | Stomach, Liver, Splenomegaly (287 mg) | / |
| KI Bcl2 Δ/+ | KI- 11 | 92 | ✓ | ✓ | Under lungs, Mesenteric LNs, Splenomegaly (145 mg) | Diffused and bloody cervical LNs |
|  | KI- 12 | 84 | X | ✓ | Splenomegaly, Liver, Kidneys | Swollen testicles |
|  | KI-13 | 117 | ✓ | ✓ | Intestine, Atrophy of all LNs, Splenomegaly (2500 mg) | / |
|  | KI-14 | 52 | X | ✓ | Atrophy of cervical LNs, Splenomegaly (300 mg) | Obese |
|  | KI-15 | 52 | X | ✓ | Splenomegaly (300 mg) | Obese |
|  | KI-16 | 41 | X | ✓ | Mesentery, Splenomegaly | / |
|  | KI-17 | 81 | X | ✓ | Mesentery, Splenomegaly | / |
|  | KI-18 | 111 | X | ✓ | Mesentery, Splenomegaly | / |
|  | KI-19 | 79 | ✓ | ✓ | Mesentery, splenomegaly (1350 mg) | Decolored liver |
| 3'RR-Bcl2+ | Tg-1 | 74 | X | ✓ | Atrophy of inguinal and axillary LNs, Splenomegaly (380) | / |
|  | Tg-2 | 74 | X | ✓ | Splenomegaly (161 mg) | / |
|  | Tg-3 | 60 | X | ✓ | Splenomegaly (240 mg) | / |
|  | Tg-4 | 60 | X | ✓ | Splenomegaly (222 mg) | / |
|  | Tg-5 | 64 | X | ✓ | Atrophy of cervical LNs<br>Splenomegaly (222 mg) | / |
|  | Tg-6 | 64 | ✓ | ✓ | Intestine, Liver, Atrophy of cervical LNs, Splenomegaly (730 mg) | / |
|  | Tg-7 | 121 | X | ✓ | Mesentery, Splenomegaly (1139 mg) | / |

**Table 2:** Table showing the detailed IgM and IgG repertoire analyses of the tumor tissues only, of a few mice from both BCL2 models. Total number of clones for each heavy chain, the corresponding number of productive reads are indicated. Based on the previous repertoire data obtained from WT mice, a certain threshold for the clone frequency was used and only the clones that were present at a frequency higher than or equal to 30% were selected. For each clone, the V and J gene used along with their corresponding V mutation frequency (without CDR3 as it is highly mutated) are also indicated.

| Genotype | Mouse nb | Heavy chain | Total number of clones | Total number of productive reads | V_gene | J_gene | Frequency (Clones ≥ 30%) | V_mutation frequency (without CDR3) 0/00 |
| --- | --- | --- | --- | --- | --- | --- | --- | --- |
| IgK-BCL2 Δ/Δ | KI-2 | **IgG** | **234** | **1269** | **IGHV4-1** | **IGHJ2** | **42.2** | **1** |
|  | KI-3 | IgM | 248 | 9087 | IGHV6-3 | IGHJ2 | 95.2 | 2.1 |
|  |  | IgG | 58 | 8308 | IGHV10-3 | IGHJ2 | 99.0 | 0.35 |
|  | KI-4 | **IgM** | **430** | **2104** | **IGHV4-1** | **IGHJ2** | **47.6** | **0** |
|  |  | **IgG** | **1055** | **6274** | **IGHV1-5** | **IGHJ3** | **30.2** | **1.5** |
|  | KI-8 | IgM | 143 | 1560 | IGHV3-6 | IGHJ2 | 64.4 | 1.8 |
|  |  | **IgG** | **26** | **2835** | **IGHV4-1** | **IGHJ2** | **98.4** | **0.8** |
| IgK-BCL2 Δ/+ | KI-16 | IgG | 15 | 871 | IGHV10-1 | IGHJ2 | 98.8 | 0 |
|  | KI-18 | IgG | 74 | 175 | IGHV10-3 | IGHJ2 | 41.1 | 0.35 |
|  | KI-19 | IgM | 494 | 2106 | IGHV1-39 | IGHJ2 | 53.4 | 0 |
|  |  | IgG | 619 | 11924 | IGHV1-26 | IGHJ2 | 85.2 | 0 |
| 3'RR-BCL2 | Tg-6 | IgG | 1157 | 12395 | IGHV2-2 | IGHJ4 | 51.1 | 0 |

*(clones carrying a VDJ N-glycosylation site are in red)*

**Figure 7:**
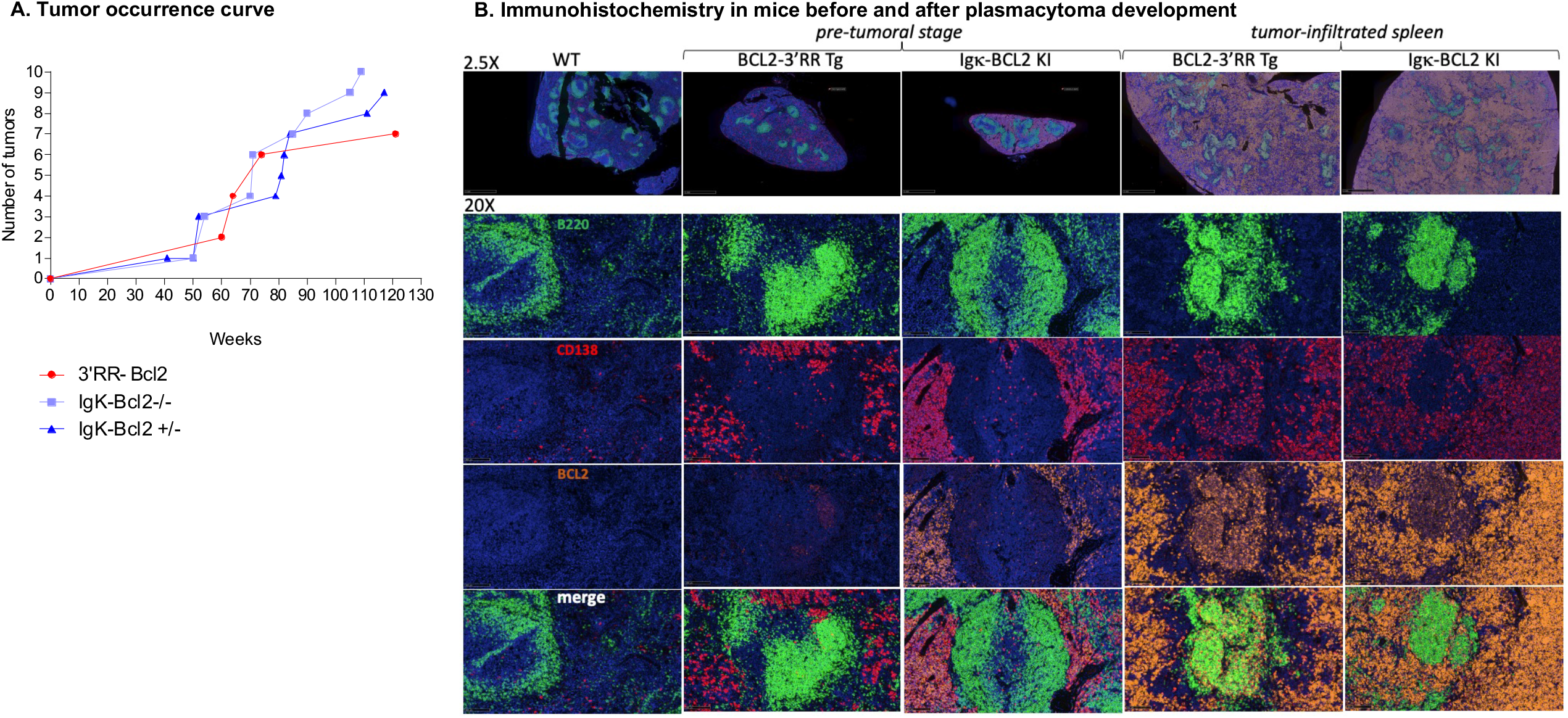
(A) Curve showing the evolution of tumor occurrence in both BCL2 models with respect to their age (in weeks). (B) Immunohistochemistry (IHC) analysis of the spleen at the pre-tumoral and tumor-infiltrated stage of both BCL2 models compared to WT. Staining was done for B cells (using B220 in Fitc → Green), plasma cells (CD138 in Tritc → Red), human BCL2 (in cyanine 5 → Yellow) and finally DAPI in blue.

Representative immunohistochemistry analyses of tissues either from young mice (thus in the pre-lymphomatous stage) or old mice (affected with overt lymphoma, notably 3’RR-*BCL2* mouse Tg-6 and Igκ-*BCL2* Δ/Δ mouse KI-8 in Table 1) are shown in Fig 6B. At the pre-lymphoma stage, the organization of the spleen is respected showing large follicles densely occupied with CD19+ B-lymphocytes, mixed with less abundant plasma cells. In such young 3’RR-BCL2 mice, BCL2 staining is mostly seen in B-cells within germinal centers. By the contrary in the Igκ-BCL2 mice, abundant plasma cells populate the red pulp already at the early stage, and stain the most strongly for BCL2.

In mice carrying tumors from either genotype, the spleen red pulp later appeared heavily infiltrated by CD138+ BCL2+ plasma cells. These tumors affected mostly the mesenteric lymph nodes, spleen and liver. The tumors analyzed at other distant locations also consisted into CD138+ cells and the disease thus appears like a disseminated plasmablastic lymphoma. (**Fig 7B**).

Ig repertoire analysis by REPseq was done for tumors from 4 Igκ-*BCL2* Δ/Δ tumoral mice (KI-2, KI-3, KI-4 and KI-8, **Table 1**), 3 Igκ-*BCL2* Δ/+ tumoral mice numbers KI-16, KI-18 and KI-19 and one 3’RR-*BCL2* tumoral mouse number Tg-6. All tumors analyzed in both strains included one or eventually two strongly predominant clonal cell populations, each so-called “predominant clonotype” representing 30% to 98% of all Ig reads (**Table 1**).

Both IgM producing and IgG clones were found, 5 of them carrying no mutation of the expressed VDJ regions, while 7 had clearly accumulated somatic hypermutation (from 3.5 to 21 / Kbp), indicating that malignant clones could originate in some cases from an extra-follicular pathway or in other cases be GC-derived.

### The *BCL2* cassette is exposed to low SHM in both transgenic models

Sequencing the promoter region of *BCL2* in both models revealed that SHM occurred all along a 1.5 kb sequence fragment, and globally appeared diversified in polyclonal cells (in agreement with the B-cell diversity indicated by Ig repertoire experiments) in mice not affected with tumors **(Fig 8**). The mutation rate at a given position thus never exceeded 0.6%. Sequences with the highest rate of SHM were rather obtained in class-switched B cells from immunized Igκ-*BCL2* mice, and the most often mutated positions were located around the P1 promoter.

**Figure 8:**
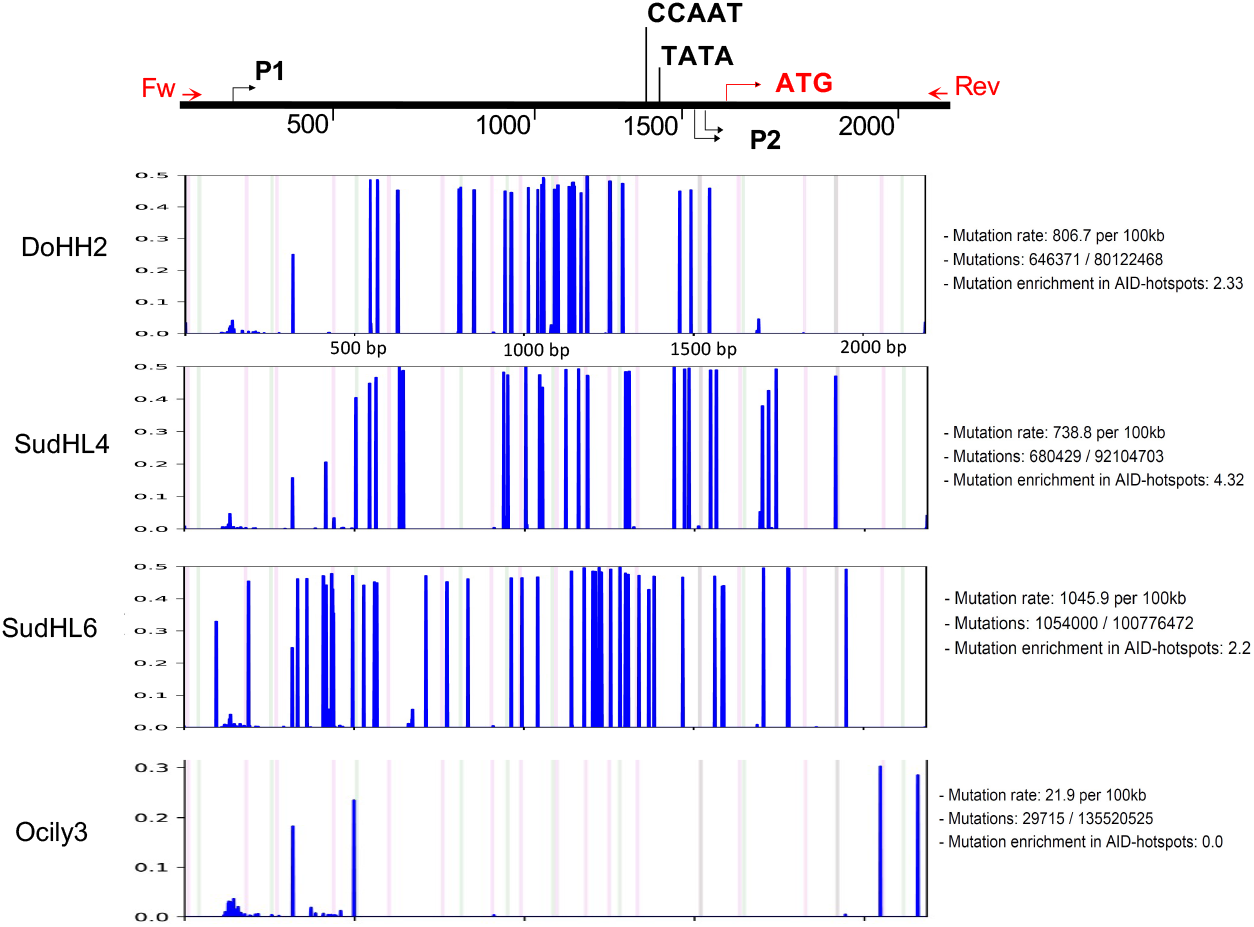
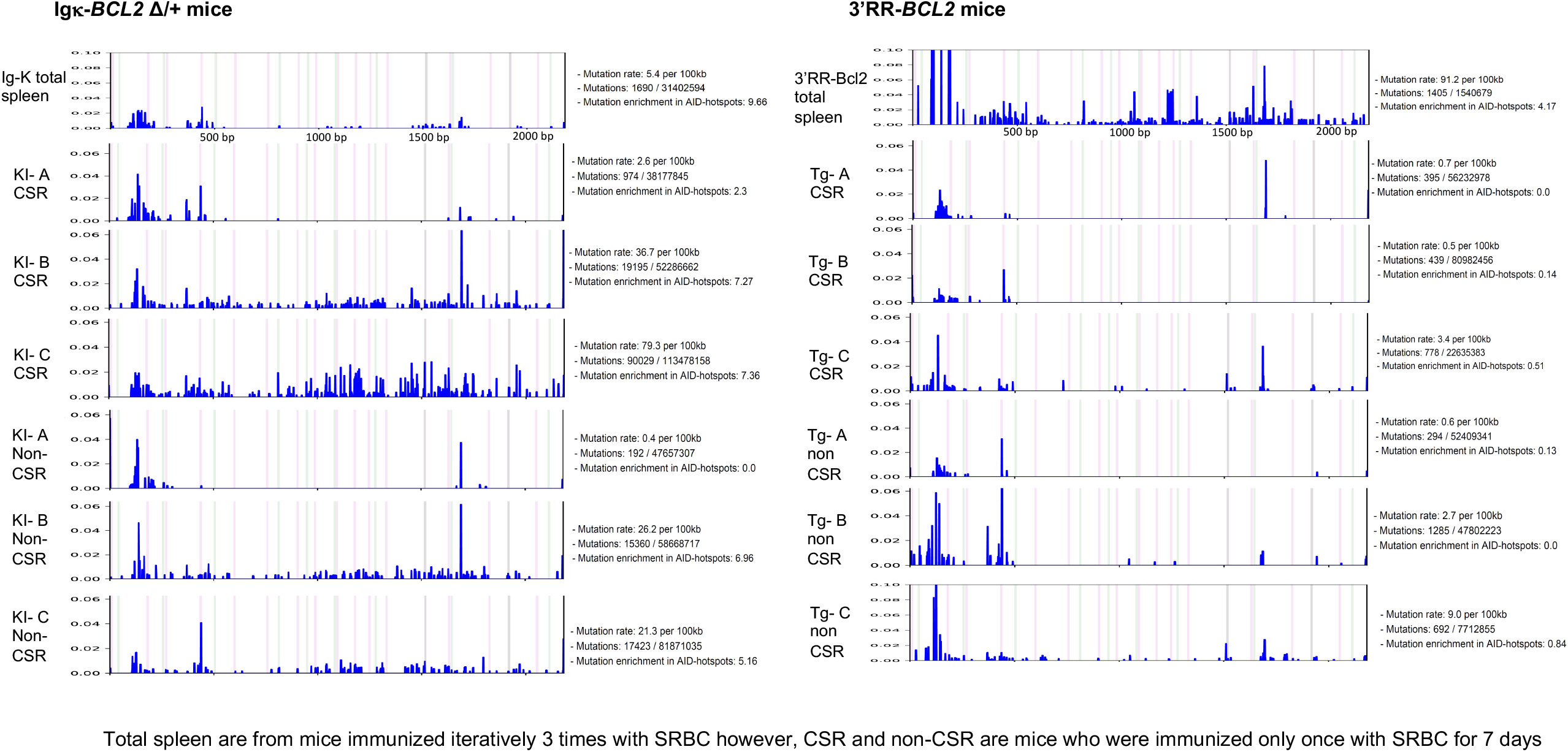
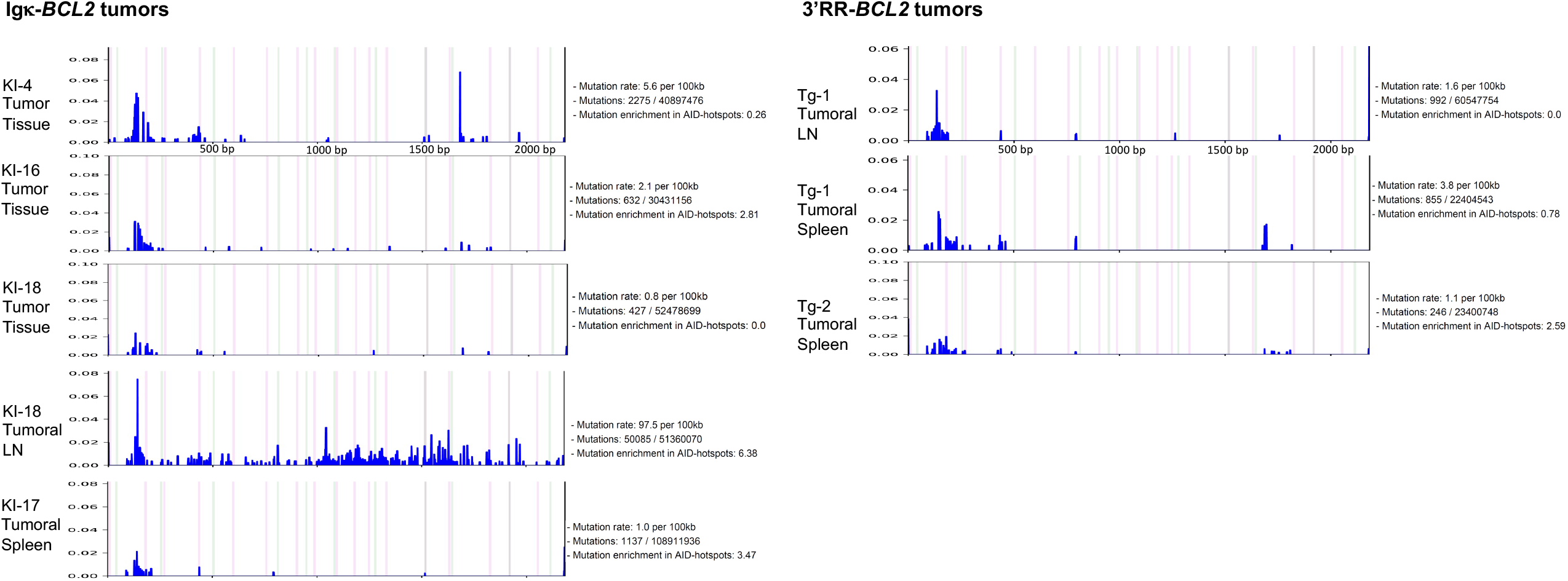
Graphs showing the frequency and the localization / distribution of mutations along the BCL2 promoter region (after high-throughput sequencing) as well as the global mutation rate and the mutation enrichment in AID-hotspots in several human DLBCL cell lines as positive controls (A), in total splenocytes or class-switched versus non-class-switched splenocytes of immunized mice (3 iterative immunizations or single 7-day immunization respectively) from both BCL2 models (B) and finally in various tissues / organs from several tumoral mice of both BCL2 models (C). Graphs are shown as generated by the system Deminer after correction with respect to two negative controls passed on the same run.

In some but not all tumors, mutations were present at a higher level, eventually approaching 10% of the reads.

## DISCUSSION

We herein report two new models of B-lineage specific BCL2 deregulation in mice. These models yield different patterns of expression, either with a global increase of BCL2 expression at all B-cell maturation stages in the Igκ-*BCL2* knock-in strain, but with a climax in plasma cells, or with a more restricted GC-specific expression in transgenics driven by the IgH locus 3’RR enhancers. Although both models roughly yield the same level of BCL2 expression in GC B cells, its higher “GC-specificity” makes the 3’RR-driven model a potentially attractive “first hit” platform for studying early events of FL lymphomagenesis. Indeed, FL is currently considered as evolving from FLLC cells carrying an initial t(14;18) translocation then followed by multiple additional driver mutations, the latter concurring to block plasma cell differentiation and exit of the transformed cells from the GC-stage.

By contrast, the strongest and broadest deregulation observed after the knock-in of BCL2 within the Igκ locus principally results into the expansion of the plasmablastic and plasma cell compartments even in young mice prior to any immunization.

Finally, regarding the occurrence of spontaneous tumors in either of these BCL2 strains, both appear exposed to plasma cell tumors, whether or not this was preceded by overt polyclonal plasmacytosis. A similar situation is likely occurring for human cells deregulating only BCL2, which circulate and can be found in blood and lymphoid tissues of many healthy individuals and do not appear to be blocked in their differentiation (Roulland et al. 2011; Sungalee et al. 2014). By contrast, local amplification of a more aggressive B-cell clone, either in the early situation of *in situ* follicular neoplasia or of overt FL is always associated to mutations additional to the BCL2 deregulation (Vogelsberg et al. 2021). The most obvious roles of all such mutations, notably involving chromatin modifier genes such as CREBBP, EZH2 or TET2, is to reprogram B-cells towards an iterative GC reentry cycle by up-regulating BCL6 expression, and inhibiting genes either involved into B-cell egress form the GC and/or into PC differentiation and progress beyond the GC B-cell stage (Lu et al. 2018; Dominguez et al. 2018; Rivas et al. 2021; Béguelin et al. 2020).

Although we have no indication about the eventual second hits supporting the growth of plasmacytoma in aged mice from this study, such acquired genetic or epigenetic anomalies are likely since no tumor arise in young mice and since, whatever the BCL2 expression dosage in both mouse strains, the clonotypic diversity of B-cells in younger tumor-free animals is normal and does not reveal the early expansion of a monoclonal population.

Contrary to our expectations, presence of a large P1/P2 BCL2 promoter fragment including the negative regulatory sites for BCL6 binding does not seem to repress expression in those mouse models, and is not affected by acquired mutations unleashing expression in some B-cell clones. It is thus likely that the spontaneous expression yielded by the BCL2 cassette in either model, is already sufficiently high by itself for initiating the clonal malignant development without the need of promoter mutations. Although the P1/P2 fragment is accessible to a significant amount of SHM, notably when inserted in the Igκ locus, these mutations rather seem to randomly accumulate at low level.

Tumors occurring in both models are thus mostly made up of clonal plasma cells (with biclonal proliferations in some cases), either producing IgG or IgM with roughly similar frequencies, and revealing SHM of VH genes ranging from 3.5 to 21 per Kbp for 7 out of 12 malignant clones, thus with a clear indication of a “post-GC” origin of the malignant clones, but without any mutation for 5 of them, which could thus likely correspond to extra-follicular PCs or to PCs differentiated independently of T-cell help. Toellner et al previously characterized extrafollicular and classical GC-derived plasma cells respectively withs means of 0.7/Kbp *vs* 5.6 Kbp (Toellner et al. 2002).

Although attributed to V(D)J recombination errors, which could potentially affect any Ig locus, BCL2 translocations associated with FL overwhelmingly involve the IgH locus on chromosome 14 and much more rarely Ig light chain loci (Szymanowska et al. 2008; Hillion et al. 1991, 2). Such translocations replace all the V constant gene cluster with BCL2 and thus associates the oncogene with regulatory elements for the constant gene cluster, *i*.*e*. predominantly under the influence of the IgH 3’RR in mature B-cells. Such conditions are thus likely to favor a strong expression in activated B-cells and at the GC-stage, as observed in our IgH-3’RR model.

Why both BCL2 strains explored in this study strongly differ for the polyclonal PC expansion developed in Igκ-BCL2 mice reveals an unexpected impact of the level of BCL2 expression into B-cell differentiation. Such an effect could either be related to an imprinting of activated B-cells, more prone to enter into the PC-differentiation pathway when BCL2 is higher, but such a differentiation bias should then decrease entry into the alternate pathway of memory B cell differentiation, which is in fact not observed. By the contrary, BCL2 transgenic B-cells show a conserved ability to mount high-affinity Ag-dependent responses after immunization. The most likely explanation for PC accumulation is thus at this stage, that the high level of BCL2 expression reached by Igκ-BCL2 cells, and to a lower extent in 3’RR-BCL2 mice, significantly impact first plasmablast, and secondarily PC survival. Initial steps of PC differentiation are indeed known to depend on endogenous BCL2 expression (Peperzak et al. 2013), and survival is thus initially increased when no decrease of BCL2 expression occurs in transgenic B cells. Later on, long-live PCs normally do not rely on BCL2 at all for their survival but rather on BCMA-dependent MCL1 expression (Peperzak et al. 2013). Since MCL1 and BCL2 share a similar pro-survival function, the knock-in Igκ-BCL2 construction maintaining or even increasing constitutive BCL2 expression in all PCs necessarily ends with deregulated survival of these cells and hypertrophy of the LLPC compartment. PCs are also strongly exposed to endothelial reticulum (ER)-mediated stress, which triggers autophagy and mitophagy. BCL2-family members such as BCL2-L13 and BNIP3 are major actors of autophagy and mitophagy (Onnis et al. 2018; Onishi et al. 2021). These factors share interactions with some BCL2 partners, such as BECLIN-1 and their function in promoting mitophagy might likely be hampered by excessive amounts of BCL2. Noticeably, human myeloma often associates up-regulation of the BCL2 homolog MCL1 and silencing of BNIP3 (Murai et al. 2005). Altogether, either by reducing ER-stress induced apoptosis, autophagy or mitophagy, the BCL2 up-regulation triggered at the plasmablast/PC stage in Igκ-BCL2 mice appears to open up the gate for entry of more B cells into the plasma cell stage together with deregulated accumulation of non-dying long-lived PCs.

## ACKNOWLEDGMENTS

This work was supported by grants from Fondation ARC pour la Recherche sur le Cancer (Grants PGA1 RF20170205386 / RF20180207070), the Institut National du cancer (INCA AAP PLBIO-17-06), the Agence Nationale de la Recherche (ANR, ANR-16-CE15-0019-01/03), the Leukemia Lymphoma Society (TRP 6593-20) and Rennes Métropole. SL was supported by a specific grant from the LabEx IGO program (n° ANR-11-LABX-0016) funded by the «Investment into the Future» French Government program, managed by the National Research Agency (ANR). The authors have no conflicting financial interests. We thank staff from the Limoges University core animal facility for animal care.

## Supplemental information

**Supplementary Table 1:** Table summarizing the list of antibodies used for the staining Panel of B and T cells in the Bone Marrow and peripheral lymphoid organs (Spleen / mesenteric Lymph nodes). Antibodies used for intracellular staining of permeabilized cells are highlighted in grey.

**Supplementary Figure 1:** (A). Gating strategy for staining of B cells in the bone marrow and periphery respectively. (B). Gating strategy for staining of B-lineage cells in the spleen or lymph nodes **(C)** Gating strategy for the staining of T cells in the periphery (staining examples are from a resting unimmunized WT mouse).

**Supplementary Figure 2**: Heatmap showing expression of the Top 10 differentially expressed genes defining the various B-cell clusters identified by single-cell transcriptome analysis.

## REFERENCES

Amin, Rada, Frédéric Mourcin, Fabrice Uhel, Céline Pangault, Philippe Ruminy, Loic Dupré, Marion Guirriec, et al. 2015. “DC-SIGN-Expressing Macrophages Trigger Activation of Mannosylated IgM B-Cell Receptor in Follicular Lymphoma.” Blood 126 (16): 1911–20. https://doi.org/10.1182/blood-2015-04-640912.

Béguelin, Wendy, Matt Teater, Cem Meydan, Kenneth B. Hoehn, Jude M. Phillip, Alexey A. Soshnev, Leandro Venturutti, et al. 2020. “Mutant EZH2 Induces a Pre-Malignant Lymphoma Niche by Reprogramming the Immune Response.” Cancer Cell 37 (5): 655–673.e11. https://doi.org/10.1016/j.ccell.2020.04.004.

Boice, Michael, Darin Salloum, Frederic Mourcin, Viraj Sanghvi, Rada Amin, Elisa Oricchio, Man Jiang, et al. 2016. “Loss of the HVEM Tumor Suppressor in Lymphoma and Restoration by Modified CAR-T Cells.” Cell 167 (2): 405–418.e13. https://doi.org/10.1016/j.cell.2016.08.032.

Butler, Andrew, Paul Hoffman, Peter Smibert, Efthymia Papalexi, and Rahul Satija. 2018. “Integrating Single-Cell Transcriptomic Data across Different Conditions, Technologies, and Species.” Nature Biotechnology 36 (5): 411–20. https://doi.org/10.1038/nbt.4096.

Chevrier, Stéphane, Céline Genton, Axel Kallies, Alexander Karnowski, Luc A. Otten, Bernard Malissen, Marie Malissen, et al. 2009. “CD93 Is Required for Maintenance of Antibody Secretion and Persistence of Plasma Cells in the Bone Marrow Niche.” Proceedings of the National Academy of Sciences of the United States of America 106 (10): 3895–3900. https://doi.org/10.1073/pnas.0809736106.

Cogné, M, R Lansford, A Bottaro, J Zhang, J Gorman, F Young, H L Cheng, and F W Alt. 1994. “A Class Switch Control Region at the 3’ End of the Immunoglobulin Heavy Chain Locus.” Cell 77 (5): 737–47.

Dominguez, Pilar M., Hussein Ghamlouch, Wojciech Rosikiewicz, Parveen Kumar, Wendy Béguelin, Lorena Fontán, Martín A. Rivas, et al. 2018. “TET2 Deficiency Causes Germinal Center Hyperplasia, Impairs Plasma Cell Differentiation, and Promotes B-Cell Lymphomagenesis.” Cancer Discovery 8 (12): 1632–53. https://doi.org/10.1158/2159-8290.CD-18-0657.

Egle, Alexander, Alan W. Harris, Mary L. Bath, Lorraine O’Reilly, and Suzanne Cory. 2004. “VavP-Bcl2 Transgenic Mice Develop Follicular Lymphoma Preceded by Germinal Center Hyperplasia.” Blood 103 (6): 2276–83. https://doi.org/10.1182/blood-2003-07-2469.

Hafemeister, Christoph, and Rahul Satija. 2019. “Normalization and Variance Stabilization of Single-Cell RNA-Seq Data Using Regularized Negative Binomial Regression.” Genome Biology 20 (1): 296. https://doi.org/10.1186/s13059-019-1874-1.

Hillion, J., C. Mecucci, A. Aventin, D. Leroux, I. Wlodarska, H. Van Den Berghe, and C. J. Larsen. 1991. “A Variant Translocation t(2;18) in Follicular Lymphoma Involves the 5’ End of Bcl-2 and Ig Kappa Light Chain Gene.” Oncogene 6 (1): 169–72.

Huet, Sarah, Pierre Sujobert, and Gilles Salles. 2018. “From Genetics to the Clinic: A Translational Perspective on Follicular Lymphoma.” Nature Reviews. Cancer 18 (4): 224–39. https://doi.org/10.1038/nrc.2017.127.

Kowalczyk, Monika S., Itay Tirosh, Dirk Heckl, Tata Nageswara Rao, Atray Dixit, Brian J. Haas, Rebekka K. Schneider, Amy J. Wagers, Benjamin L. Ebert, and Aviv Regev. 2015. “Single-Cell RNA-Seq Reveals Changes in Cell Cycle and Differentiation Programs upon Aging of Hematopoietic Stem Cells.” Genome Research 25 (12): 1860–72. https://doi.org/10.1101/gr.192237.115.

Lin, Pei, Rechna Jetly, Patrick A. Lennon, Lynne V. Abruzzo, Sapana Prajapati, and L. Jeffrey Medeiros. 2008. “Translocation (18;22)(Q21;Q11) in B-Cell Lymphomas: A Report of 4 Cases and Review of the Literature.” Human Pathology 39 (11): 1664–72. https://doi.org/10.1016/j.humpath.2008.04.007.

Lu, Xiaoqing, Tharu M. Fernando, Chen Lossos, Nevin Yusufova, Fan Liu, Lorena Fontán, Matthew Durant, et al. 2018. “PRMT5 Interacts with the BCL6 Oncoprotein and Is Required for Germinal Center Formation and Lymphoma Cell Survival.” Blood 132 (19): 2026–39. https://doi.org/10.1182/blood-2018-02-831438.

Martin, Ophélie Alyssa, Armand Garot, Sandrine Le Noir, Jean-Claude Aldigier, Michel Cogné, Eric Pinaud, and François Boyer. 2018. “Detecting Rare AID-Induced Mutations in B-Lineage Oncogenes from High-Throughput Sequencing Data Using the Detection of Minor Variants by Error Correction Method.” The Journal of Immunology 201 (3): 950–56. https://doi.org/10.4049/jimmunol.1800203.

Mayer, Christian T., Anna Gazumyan, Ervin E. Kara, Alexander D. Gitlin, Jovana Golijanin, Charlotte Viant, Joy Pai, et al. 2017. “The Microanatomic Segregation of Selection by Apoptosis in the Germinal Center.” Science (New York, N.Y.) 358 (6360). https://doi.org/10.1126/science.aao2602.

McDonnell, T. J., N. Deane, F. M. Platt, G. Nunez, U. Jaeger, J. P. McKearn, and S. J. Korsmeyer. 1989. “Bcl-2-Immunoglobulin Transgenic Mice Demonstrate Extended B Cell Survival and Follicular Lymphoproliferation.” Cell 57 (1): 79–88. https://doi.org/10.1016/0092-8674(89)90174-8.

McDonnell, T. J., and S. J. Korsmeyer. 1991. “Progression from Lymphoid Hyperplasia to High-Grade Malignant Lymphoma in Mice Transgenic for the t(14; 18).” Nature 349 (6306): 254–56. https://doi.org/10.1038/349254a0.

Milpied, Pierre, Anita K. Gandhi, Guillaume Cartron, Laura Pasqualucci, Karin Tarte, Bertrand Nadel, and Sandrine Roulland. 2021. “Follicular Lymphoma Dynamics.” Advances in Immunology 150: 43–103. https://doi.org/10.1016/bs.ai.2021.05.002.

Milpied, Pierre, Bertrand Nadel, and Sandrine Roulland. 2015. “Premalignant Cell Dynamics in Indolent B-Cell Malignancies.” Current Opinion in Hematology 22 (4): 388–96. https://doi.org/10.1097/MOH.0000000000000159.

Monni, O., H. Joensuu, K. Franssila, J. Klefstrom, K. Alitalo, and S. Knuutila. 1997. “BCL2 Overexpression Associated with Chromosomal Amplification in Diffuse Large B-Cell Lymphoma.” Blood 90 (3): 1168–74.

Murai, M., M. Toyota, A. Satoh, H. Suzuki, K. Akino, H. Mita, Y. Sasaki, et al. 2005. “Aberrant DNA Methylation Associated with Silencing BNIP3 Gene Expression in Haematopoietic Tumours.” British Journal of Cancer 92 (6): 1165–72. https://doi.org/10.1038/sj.bjc.6602422.

Ogilvy, S., D. Metcalf, C. G. Print, M. L. Bath, A. W. Harris, and J. M. Adams. 1999. “Constitutive Bcl-2 Expression throughout the Hematopoietic Compartment Affects Multiple Lineages and Enhances Progenitor Cell Survival.” Proceedings of the National Academy of Sciences of the United States of America 96 (26): 14943–48. https://doi.org/10.1073/pnas.96.26.14943.

Onishi, Mashun, Koji Yamano, Miyuki Sato, Noriyuki Matsuda, and Koji Okamoto. 2021. “Molecular Mechanisms and Physiological Functions of Mitophagy.” The EMBO Journal 40 (3): e104705. https://doi.org/10.15252/embj.2020104705.

Onnis, Anna, Valentina Cianfanelli, Chiara Cassioli, Dijana Samardzic, Pier Giuseppe Pelicci, Francesco Cecconi, and Cosima T. Baldari. 2018. “The Pro-Oxidant Adaptor P66SHC Promotes B Cell Mitophagy by Disrupting Mitochondrial Integrity and Recruiting LC3-II.” Autophagy 14 (12): 2117–38. https://doi.org/10.1080/15548627.2018.1505153.

Peperzak, Victor, Ingela Vikström, Jennifer Walker, Stefan P. Glaser, Melanie LePage, Christine M. Coquery, Loren D. Erickson, et al. 2013. “Mcl-1 Is Essential for the Survival of Plasma Cells.” Nature Immunology 14 (3): 290–97. https://doi.org/10.1038/ni.2527.

Péron, Sophie, Brice Laffleur, Nicolas Denis-Lagache, Jeanne Cook-Moreau, Aurélien Tinguely, Laurent Delpy, Yves Denizot, Eric Pinaud, and Michel Cogné. 2012. “AID-Driven Deletion Causes Immunoglobulin Heavy Chain Locus Suicide Recombination in B Cells.” Science (New York, N.Y.) 336 (6083): 931–34. https://doi.org/10.1126/science.1218692.

Pinaud, Eric, Marie Marquet, Rémi Fiancette, Sophie Péron, Christelle Vincent-Fabert, Yves Denizot, and Michel Cogné. 2011. “The IgH Locus 3’ Regulatory Region: Pulling the Strings from Behind.” Advances in Immunology 110: 27–70. https://doi.org/10.1016/B978-0-12-387663-8.00002-8.

Raghavan, Sathees C., Patrick C. Swanson, Xiantuo Wu, Chih-Lin Hsieh, and Michael R. Lieber. 2004. “A Non-B-DNA Structure at the Bcl-2 Major Breakpoint Region Is Cleaved by the RAG Complex.” Nature 428 (6978): 88–93. https://doi.org/10.1038/nature02355.

Rivas, Martín A., Cem Meydan, Christopher R. Chin, Matt F. Challman, Daleum Kim, Bhavneet Bhinder, Andreas Kloetgen, et al. 2021. “Smc3 Dosage Regulates B Cell Transit through Germinal Centers and Restricts Their Malignant Transformation.” Nature Immunology 22 (2): 240–53. https://doi.org/10.1038/s41590-020-00827-8.

Rouaud, Pauline, Christelle Vincent-Fabert, Alexis Saintamand, Rémi Fiancette, Marie Marquet, Isabelle Robert, Bernardo Reina-San-Martin, Eric Pinaud, Michel Cogné, and Yves Denizot. 2013. “The IgH 3’ Regulatory Region Controls Somatic Hypermutation in Germinal Center B Cells.” The Journal of Experimental Medicine 210 (8): 1501–7. https://doi.org/10.1084/jem.20130072.

Roulland, Sandrine, Mustapha Faroudi, Emilie Mamessier, Stéphanie Sungalee, Gilles Salles, and Bertrand Nadel. 2011. “Early Steps of Follicular Lymphoma Pathogenesis.” Advances in Immunology 111: 1–46. https://doi.org/10.1016/B978-0-12-385991-4.00001-5.

Ruminy, P., F. Jardin, J. M. Picquenot, P. Gaulard, F. Parmentier, G. Buchonnet, C. Maisonneuve, H. Tilly, and C. Bastard. 2006. “Two Patterns of Chromosomal Breakpoint Locations on the Immunoglobulin Heavy-Chain Locus in B-Cell Lymphomas with t(3;14)(Q27;Q32): Relevance to Histology.” Oncogene 25 (35): 4947–54. https://doi.org/10.1038/sj.onc.1209512.

Saintamand, Alexis, Pauline Rouaud, Faten Saad, Géraldine Rios, Michel Cogné, and Yves Denizot. 2015. “Elucidation of IgH 3’ Region Regulatory Role during Class Switch Recombination via Germline Deletion.” Nature Communications 6: 7084. https://doi.org/10.1038/ncomms8084.

Saito, Masumichi, Urban Novak, Erich Piovan, Katia Basso, Pavel Sumazin, Christof Schneider, Marta Crespo, et al. 2009. “BCL6 Suppression of BCL2 via Miz1 and Its Disruption in Diffuse Large B Cell Lymphoma.” Proceedings of the National Academy of Sciences of the United States of America 106 (27): 11294–99. https://doi.org/10.1073/pnas.0903854106.

Smith, K. G., A. Light, L. A. O’Reilly, S. M. Ang, A. Strasser, and D. Tarlinton. 2000. “Bcl-2 Transgene Expression Inhibits Apoptosis in the Germinal Center and Reveals Differences in the Selection of Memory B Cells and Bone Marrow Antibody-Forming Cells.” The Journal of Experimental Medicine 191 (3): 475–84. https://doi.org/10.1084/jem.191.3.475.

Strasser, A., A. W. Harris, M. L. Bath, and S. Cory. 1990. “Novel Primitive Lymphoid Tumours Induced in Transgenic Mice by Cooperation between Myc and Bcl-2.” Nature 348 (6299): 331–33. https://doi.org/10.1038/348331a0.

Strasser, A., S. Whittingham, D. L. Vaux, M. L. Bath, J. M. Adams, S. Cory, and A. W. Harris. 1991. “Enforced BCL2 Expression in B-Lymphoid Cells Prolongs Antibody Responses and Elicits Autoimmune Disease.” Proceedings of the National Academy of Sciences of the United States of America 88 (19): 8661–65.

Stuart, Tim, Andrew Butler, Paul Hoffman, Christoph Hafemeister, Efthymia Papalexi, William M. Mauck, Yuhan Hao, Marlon Stoeckius, Peter Smibert, and Rahul Satija. 2019. “Comprehensive Integration of Single-Cell Data.” Cell 177 (7): 1888–1902.e21. https://doi.org/10.1016/j.cell.2019.05.031.

Sungalee, Stéphanie, Emilie Mamessier, Ester Morgado, Emilie Grégoire, Philip Z. Brohawn, Christopher A. Morehouse, Nathalie Jouve, et al. 2014. “Germinal Center Reentries of BCL2-Overexpressing B Cells Drive Follicular Lymphoma Progression.” The Journal of Clinical Investigation 124 (12): 5337–51. https://doi.org/10.1172/JCI72415.

Szymanowska, Natalia, Wolfram Klapper, Stefan Gesk, Ralf Küppers, José I. Martín-Subero, and Reiner Siebert. 2008. “BCL2 and BCL3 Are Recurrent Translocation Partners of the IGH Locus.” Cancer Genetics and Cytogenetics 186 (2): 110–14. https://doi.org/10.1016/j.cancergencyto.2008.06.007.

Tellier, Julie, Cedric Menard, Sandrine Roulland, Nadine Martin, Céline Monvoisin, Lionel Chasson, Bertrand Nadel, Philippe Gaulard, Claudine Schiff, and Karin Tarte. 2014. “Human t(14;18)Positive Germinal Center B Cells: A New Step in Follicular Lymphoma Pathogenesis?” Blood 123 (22): 3462–65. https://doi.org/10.1182/blood-2013-12-545954.

Toellner, Kai-Michael, William E. Jenkinson, Dale R. Taylor, Mahmood Khan, Daniel M.-Y. Sze, David M. Sansom, Carola G. Vinuesa, and Ian C. M. MacLennan. 2002. “Low-Level Hypermutation in T Cell-Independent Germinal Centers Compared with High Mutation Rates Associated with T Cell-Dependent Germinal Centers.” The Journal of Experimental Medicine 195 (3): 383–89. https://doi.org/10.1084/jem.20011112.

Vogelsberg, Antonio, Julia Steinhilber, Barbara Mankel, Birgit Federmann, Janine Schmidt, Ivonne A. Montes-Mojarro, Katrin Hüttl, et al. 2021. “Genetic Evolution of in Situ Follicular Neoplasia to Aggressive B-Cell Lymphoma of Germinal Center Subtype.” Haematologica 106 (10): 2673–81. https://doi.org/10.3324/haematol.2020.254854.

Zhang, Chunmei, Rende Li, Yongming Li, Chaojun Song, Zhijia Liu, Yun Zhang, Zhuwei Xu, et al. 2012. “Establishment of Reverse Direct ELISA and Its Application in Screening High-Affinity Monoclonal Antibodies.” Hybridoma 31 (4): 284–88. https://doi.org/10.1089/hyb.2012.0010.

