## Supplemental Table and figures for "Diverse B-cell specific transcriptional contexts of the BCL2 oncogene in mouse models impacts pre-malignant development"

**Supplemental information**

**Supplementary Table 1:** Table summarizing the list of antibodies used for the staining Panel of B and T cells in the Bone Marrow and peripheral lymphoid organs (Spleen / mesenteric Lymph nodes). Antibodies used for intracellular staining of permeabolized cells are highlighted in grey.

**Supplementary Figure 1:** (A). Gating strategy for staining of B cells in the bone marrow and periphery respectively. (B). Gating strategy for staining of B-lineage cells in the spleen or lymph nodes **(C)** Gating strategy for the staining of T cells in the periphery (staining examples are from a resting unimmunized WT mouse).

**Supplementary Figure 2**: Heatmap showing expression of the Top 10 differentially expressed genes defining the various B-cell clusters identified by single-cell transcriptome analysis.

Supplementary Table 1: Staining Panel

| LB Moelle (BM) | | | | Peripheral B cells (Spleen + mesenteric Lymph nodes) | | | | T cells (Spleen + mesenteric Lymph nodes) | | | |
|---|---|---|---|---|---|---|---|---|---|---|---|
| Antibody | Reference | Supplier | Dilution | Antibody | Reference | Supplier | Dilution | Antibody | Reference | Supplier | Dilution |
| Bcl2 BV421 | Bcl-2/100 | BD Biosciences | 1/50 | Bcl2 BV421 | Bcl-2/100 | BD Biosciences | 1/50 | Ki-67 BV786 | B56 | BD Biosciences | 1/50 |
| CD19 BV510 | 1D3 | BD Biosciences | 1/200 | CD19 BV510 | 1D3 | BD Biosciences | 1/200 | Bcl6 FITC | K112-91 | BD Biosciences | 1/50 |
| CD24 BV650 | M1/69 | BD Biosciences | 1/400 | CD93 BV650 | AA4.1 | BD Biosciences | 1/200 | fox P3 PE | FJK-16s | Invitrogen | 1/100 |
| CD23 BV711 | B3B4 | BD Biosciences | 1/100 | CD138 Biotin | 281-2 | Biolegend | 1/200 | CD62L BV421 | MEL-14 | BD Biosciences | 1/1000 |
| B220Bv786 | RA3-6B2 | BD Biosciences | 1/100 | CD23 FITC/ A488 | B3B4 | Biolegend | 1/100 | CD4 PE-CF594 | RM4-5 | BD Biosciences | 1/400 |
| CD2 FITC/A488 | RM2-5 | BD Biosciences | 1/800 | IgD PerCP-Cy5.5 | 11-26c.2a | BD Biosciences | 1/100 | Icos PerCP-Cy5.5 | C398.4A | Biolegend | 1/100 |
| CD117 PerCP-Cy5.5 | 2B8 | Biolegend | 1/50 | CXCR4 PE | 551966 | BD Biosciences | 1/50 | PD1 PE-Cy7 | 29F.1A12 | Biolegend | 1/50 |
| CD43 PE | S7 | BD Biosciences | 1/50 | CD21 PE-CF594 | 7G6 | BD Biosciences | 1/400 | CXCR5 APC | REA215 | Milteny Biotech | 1/50 |
| BP1 Biotin | 6C3 | BD Biosciences | 1/25 | IgM PC7 | eB121-15F9 | Invitrogen | 1/200 | CD44 APC-cy7 | IM7 | BD Biosciences | 1/100 |
| IgM PE-Cy5 | II/41 | eBiosciences | 1/100 | Gl7 APC | GL7 | Invitrogen | 1/100 | FVS 510 | 564406 | BD Biosciences | 1/1000 |
| CD25 PE-Cy7 | PC61 | Biolegend | 1/50 | CD38 APC-R700 | 90 | Invitrogen | 1/100 | | | | |
| CD127 APC | 564175 | BD Biosciences | 1/50 | Streptavidin BV786 | 563858 | BD Biosciences | 1/400 | | | | |
| CD138 APC-R700 | 281-2 | BD Biosciences | 1/200 | FVS 780 APC-H7 | 565388 | BD Biosciences | 1/1000 | | | | |
| Streptavidin PE-CF594 | 562318 | BD Biosciences | 1/400 | | | | | | | | |
| FVS 780 APC-H7 | 565388 | BD Biosciences | 1/1000 | | | | | | | | |

Antibodies used for intracelllar staining of permeabilized cells are highlighted in Grey

Supplementary Figure 1A : Gating strategy for B cells in the Bone marrow

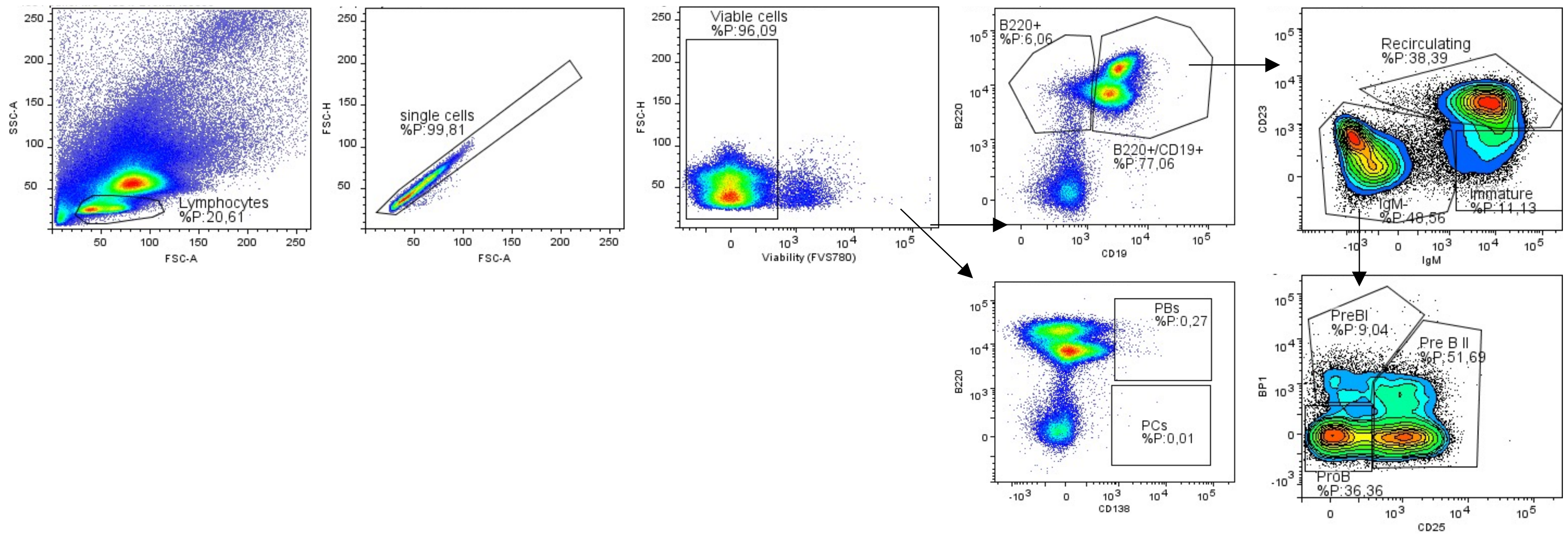

Supplementary Figure 1B: Gating strategy for peripheral B cells (spleen and mesenteric LNs)

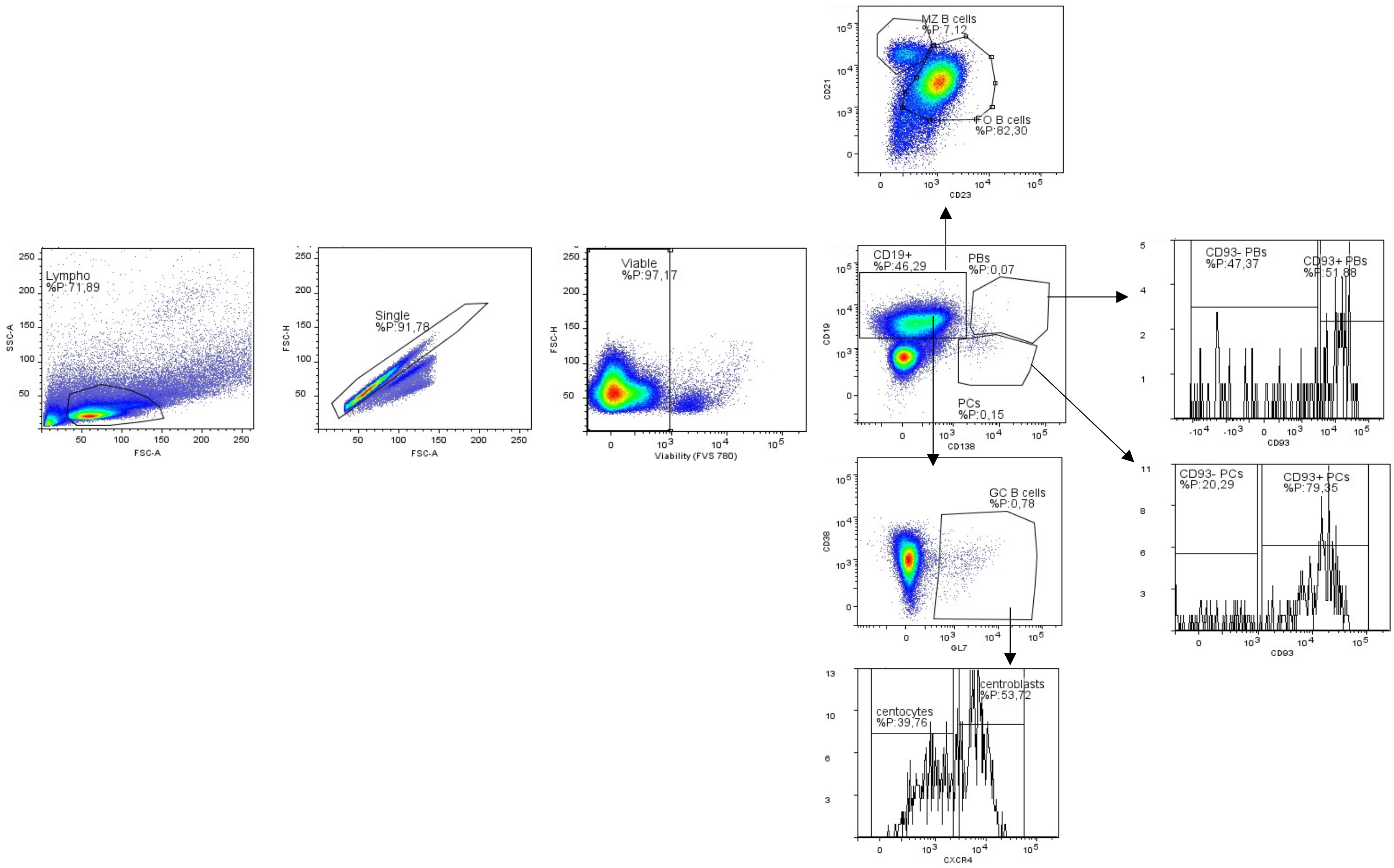

Supplementary Figure 1C: Gating strategy for peripheral T cells (in spleen and mesenteric LNs)

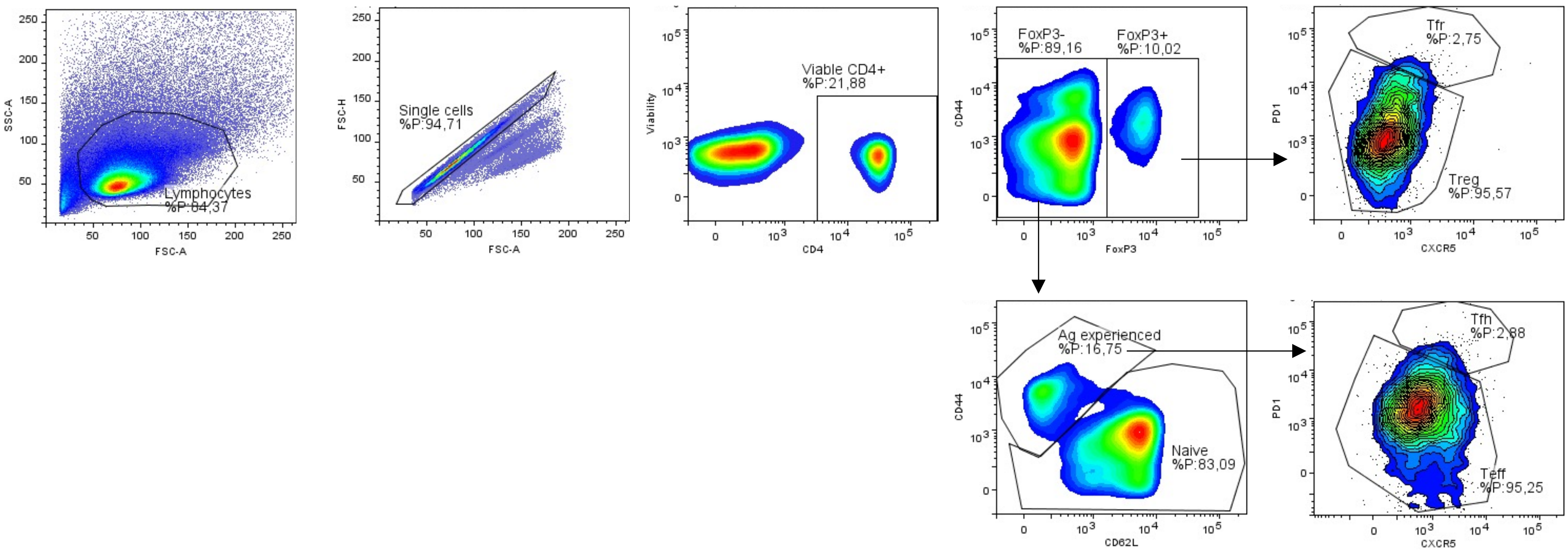

Supplementary Figure 2: Top 10 differentially expressed genes in clusters.

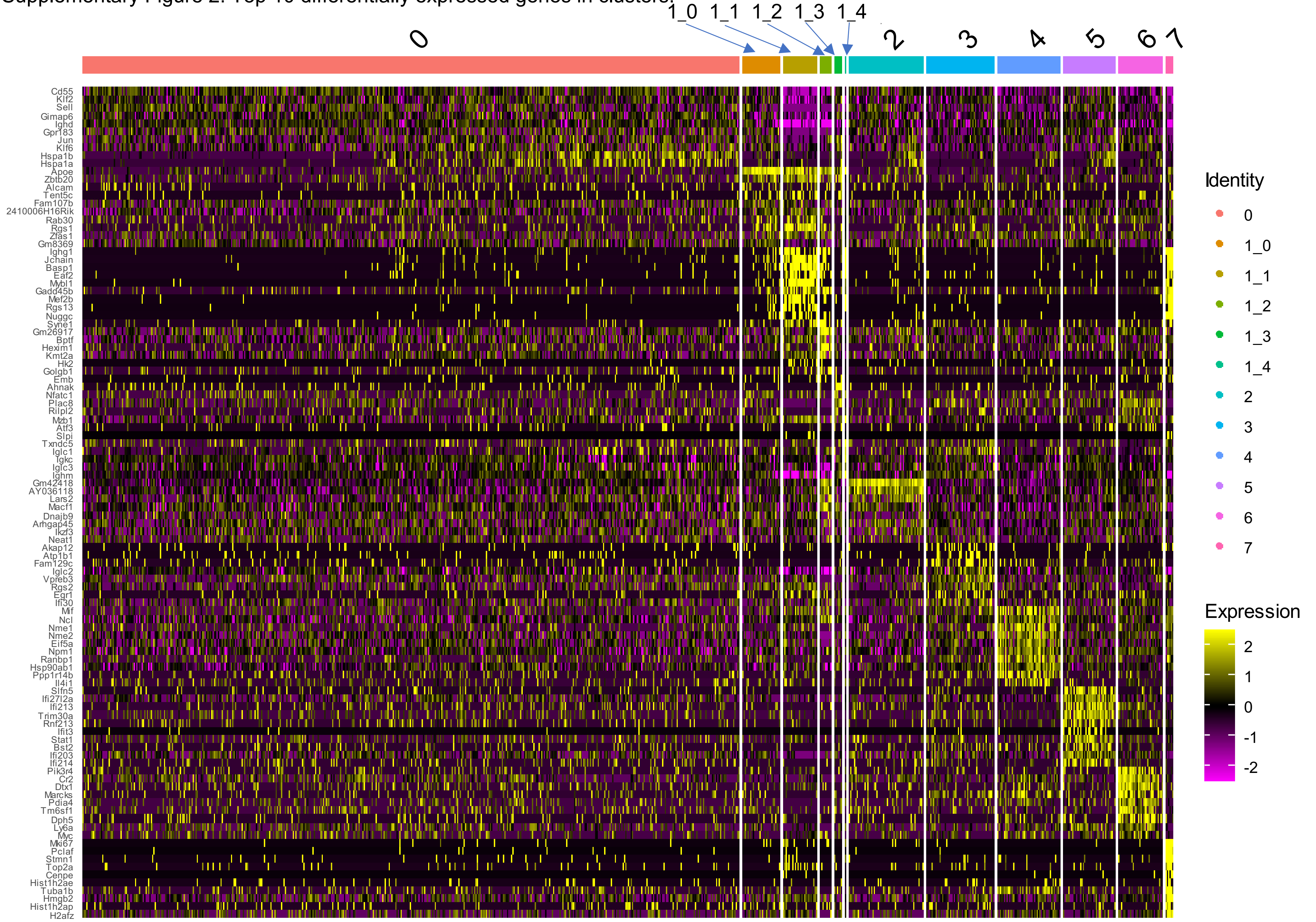
